# Sequence, Structure and Context Preferences of Human RNA Binding Proteins

**DOI:** 10.1101/201996

**Authors:** Daniel Dominguez, Peter Freese, Maria Alexis, Amanda Su, Myles Hochman, Tsultrim Palden, Cassandra Bazile, Nicole J Lambert, Eric L Van Nostrand, Gabriel A. Pratt, Gene W. Yeo, Brenton R. Graveley, Christopher B. Burge

## Abstract

Production of functional cellular RNAs involves multiple processing and regulatory steps principally mediated by RNA binding proteins (RBPs). Here we present the affinity landscapes of 78 human RBPs using an unbiased assay that determines the sequence, structure, and context preferences of an RBP *in vitro* from deep sequencing of bound RNAs. Analyses of these data revealed several interesting patterns, including unexpectedly low diversity of RNA motifs, implying frequent convergent evolution of binding specificity toward a relatively small set of RNA motifs, many with low compositional complexity. Offsetting this trend, we observed extensive preferences for contextual features outside of core RNA motifs, including spaced “bipartite” motifs, biased flanking nucleotide context, and bias away from or towards RNA structure. These contextual features are likely to enable targeting of distinct subsets of transcripts by different RBPs that recognize the same core motif. Our results enable construction of “RNA maps” of RBP activity without requiring crosslinking-based assays, and provide unprecedented depth of information on the interaction of RBPs with RNA.

## INTRODUCTION

RNA binding proteins (RBPs) control the production, maturation, localization, modification, translation, and degradation of cellular RNAs. Many RBPs contain well-defined RNA binding domains (RBDs) that engage RNA in a sequence-and/or structure-specific manner. The human genome encodes at least 1500 RBPs that contain established RBDs, the most prevalent of which include RNA recognition motifs (RRM, ~240 RBPs), HNRNP K-homology domains (KH, ~60 RBPs) and C3H1 zinc-finger domains (ZNFs, ~50 RBPs) (reviewed by (Gerstberger et al., 2014)). While RBPs containing RRM (Query et al., 1989) or KH domains (Siomi et al., 1993) were first described over two decades ago, the repertoire of RNA sequences and cellular targets bound by different members of these and other classes of RBPs are still largely unknown.

Structural studies have revealed conserved residues that enable canonical RBP-RNA interactions but have also uncovered non-canonical binding modes that have made it difficult to infer RNA target preferences from amino acid sequence alone (reviewed by (Cléry and Allain, 2013; Valverde et al., 2008)). For example, RRMs adopt a structure with an antiparallel four-stranded beta sheet packed onto two alpha helices, with the two central strands (RNP1 and RNP2) typically mediating interactions required for binding (reviewed by (Afroz et al., 2015)). However, crystallographic and NMR-based studies have shown that certain RBPs bind RNA via the linker regions, loops, or the C-and N-terminal extremities of their RRMs rather than the canonical RNP1 and RNP2 strands (reviewed by (Daubner et al., 2013)). Similarly, KH domains form a hydrophobic binding cleft that is generally thought to accommodate a pyrimidine-rich tetranucleotide motif, but specificity is often modulated by hydrogen bonding or additional interactions with the protein backbone (Grishin, 2001; Valverde et al., 2008). These variable RNA binding mechanisms in combination with the presence of multiple RBDs in most RBPs (Lunde et al., 2007) have motivated efforts to deeply interrogate the specificity of individual RBPs (reviewed by (Cléry and Allain, 2013)).

Several methods exist for determining RBP binding sites *in vivo,* most notably RNA immunoprecipitation (RIP) and UV crosslinking followed by immunoprecipitation (CLIP), followed by sequencing (Gilbert and Svejstrup, 2006; Ule et al., 2003). While such techniques capture RBP-RNA interactions in their cellular contexts, it is often difficult to derive motifs from these experiments due to interactions with protein cofactors, high levels of non-specific background (Friedersdorf and Keene, 2014), and non-random genomic composition. Quantitative *in vitro* assays such as electrophoretic mobility shift assay (EMSA), surface plasmon resonance (SPR), and isothermal calorimetry (ITC) must be guided by *a priori* knowledge of putative RNA substrates, making them unsuitable for high-throughput motif discovery. Methods such as SELEX (systematic evolution of ligands by exponential selection) typically select for a few high-affinity ‘winner’ sequences *de novo*, but generally do not reveal the full spectrum of RNA targets or their associated affinities (reviewed by (Cook et al., 2015)). RNAcompete is a high-throughput *in vitro* binding assay that captures a more complete specificity profile by quantifying the relative affinity of an RBP for a pre-defined set of ~250,000 RNA molecules (Ray et al., 2009). One limitation of this approach is that the designed RNAs present motifs in a relatively small range of predominantly unstructured contexts, restricting the analysis to short, mostly unpaired motifs. More recent approaches such as RNA Bind-n-Seq (Lambert et al., 2014) and RNAcompeteS (Cook et al., 2017) perform high-throughput sequencing of bound RNAs selected from a random pool, yielding a more comprehensive profile of the sequence and RNA secondary structural specificity of an RBP.

One or more of the *de novo* methods described above have been used to characterize the specificity of ~100 human RBPs (Giudice et al., 2016). However, integrated analysis and comparison of their specificities is made difficult by the diversity of techniques used. To systematically explore the spectrum of RNA binding specificities represented by the human proteome at high resolution, we performed RBNS on a diverse set of 78 human RBPs, half of which had previously uncharacterized specificities. RBNS comprehensively and quantitatively maps the RNA binding specificity spectrum of an RBP using a one-step *in vitro* binding reaction using recombinant RBP incubated with a random pool of RNA oligonucleotides (Lambert et al., 2014). The assay was carried out for each RBP at five protein concentrations, for a total of 400 binding assays yielding over 6 billion protein-associated reads, which enabled detection not only of simple sequence motifs but also of preferred structural and contextual features (**Fig. 1A**). Analysis of sequence motifs yielded a surprising degree of overlap in the motifs recognized by unrelated RBPs, and a bias for low-complexity motifs. However, analysis of contextual features using custom software pipelines revealed extensive divergence among proteins with similar primary motifs in additional binding features such as RNA secondary structure, flanking nucleotide context, and bipartite motifs. Thus, the diversity of regulatory targets recognized by human RBPs results from extensive reliance on features outside of core motifs.

**Figure 1.**
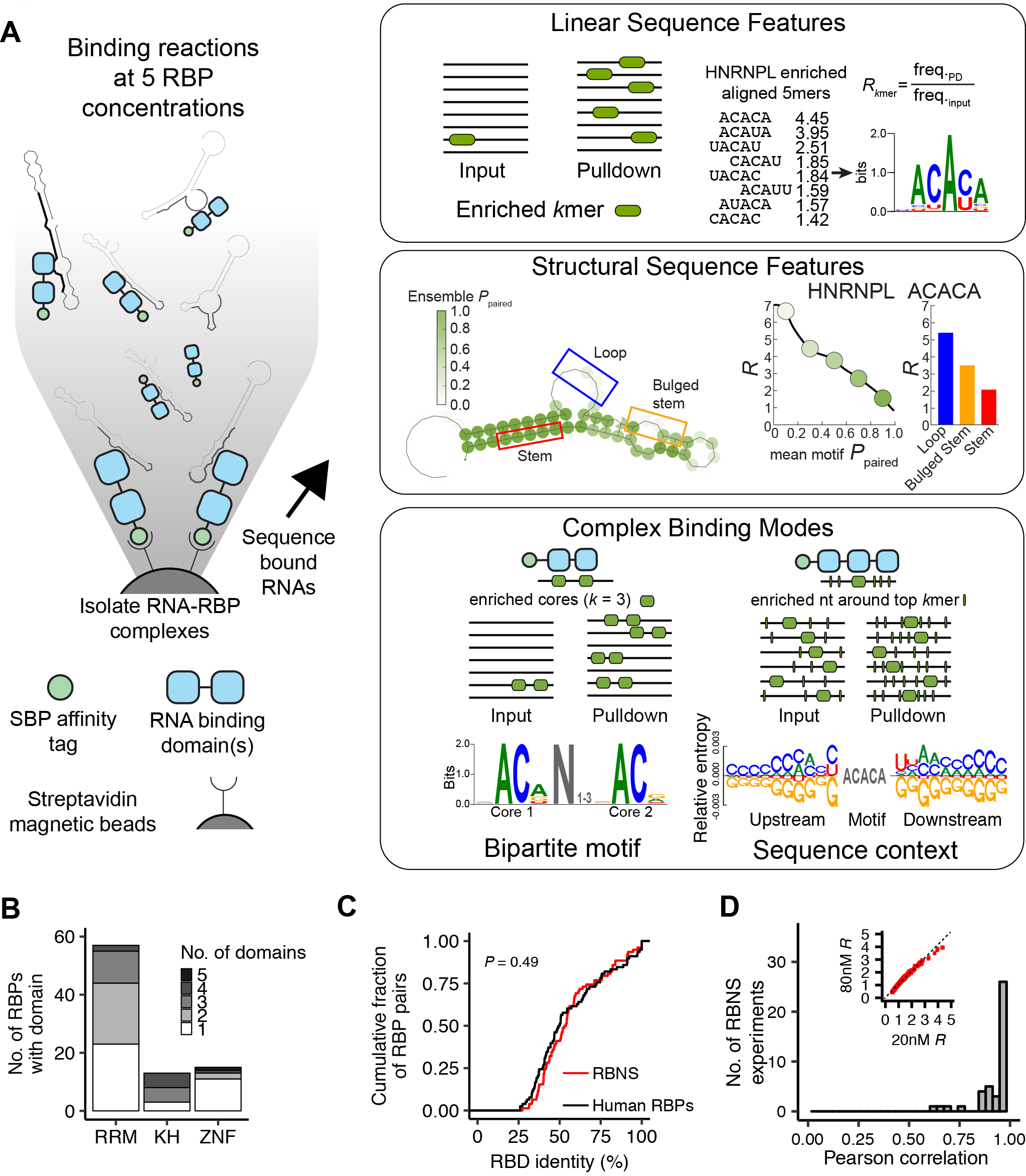
Overview of the high-throughput RNA Bind-n-Seq assay and computational analysis pipeline. **A.** Schematic of RBNS assay and pipeline. Recombinant RBPs are incubated with a pool of random RNA (black) flanked by adaptor sequences (gray). RBP-RNA complexes are isolated with streptavadin magnetic beads and bound RNA is sequenced. Computational analysis of pulldown and input reads reveals linear sequence specificities, secondary structure preferences, and complex binding modes of RBPs. **B.** Number of RBPs with one or more of the three most common RBD types assayed. **C.** Cumulative distribution of the RBD identity between each RBP and its most similar RBNS RBP. Distributions are calculated separately for the set of RBPs that has been assayed by RBNS and all other human RBPs (sampled to match RBNS domain distributions). Only domains with 5+ RBPs assayed by RBNS are included (RRM, KH, Zinc finger CCCH-type). **D.** Histogram of Pearson correlations between RBNS assays of the same RBP at different protein concentrations. Inset: correlation of 5mer *R* values of HNRNPL at 20 nM (most enriched concentration) and 80 nM.

## RESULTS

### High-throughput RNA Bind-n-Seq Assay

To determine the detailed binding preferences of a large set of human RBPs we developed a high-throughput version of RNA Bind-n-Seq (RBNS), an *in vitro* method capable of determining the sequence, structure and context preferences of RBPs. In this assay, randomized RNA oligonucleotides (20 or 40 nt) flanked by constant adapter sequences were synthesized and incubated with varying concentrations of an SBP-tagged recombinant protein containing the RBD(s) of an RBP (**Fig. 1A**, constructs listed in **Table S2**). RNA-protein complexes were isolated with streptavidin-conjugated affinity resin, washed, and bound RNA was eluted and prepared for deep sequencing. Protein purification, binding assays, and sequencing library preparations were carried out in 96-well format increasing scalability and consistency across experiments (**Methods**). A typical experiment yielded ~10-20 million unique reads at each protein concentration, which were compared to the input RNA pool sequenced to similar depth (**Fig. S1A**, **Table S2**). Inclusion of sequencing adapters flanking the randomized RNA region simplified library preparation, bypassing ligation steps which can introduce biases and preventing amplification of contaminating bacterial RNA carried over from protein purification (Lambert et al., 2014). Furthermore, as RBPs bind RNA motifs in a wide range of specific structural contexts *in vivo* (Fukunaga et al., 2014), RBNS presents RBPs with motifs spanning a broad spectrum of secondary structures, exceeding that of similar reported methods (Cook et al., 2015) (**Fig. S1B**). The high sequence complexity of the interrogated libraries enabled the fine dissection of RNA binding preferences, and use of multiple protein concentrations increased reliability and enabled detection of lower-affinity motifs.

### Binding specificities of a diverse set of human RNA binding proteins

RBNS was performed on a fairly diverse set of 78 human RBPs containing a variety of types and numbers of RBDs (**Fig. 1B**). The set of RBPs was chosen based on a combination of criteria, including: presence of well-established RBDs; evidence of role in RNA biology (though this was not required); and secondary criteria related to expression in ENCODE cell lines K562 and HepG2 and availability of validated antibodies for complementary eCLIP analysis (Sundararaman et al., 2016). Comparing all analyzed RBDs, the range of amino acid identity was similar to that of human RBPs overall (**Fig. 1C**). Together, this set captures a broad swath of proteins that is reasonably representative of human RBPs overall.

To assess the sequence specificity of each RBP, we developed a computational pipeline that calculates enrichment (“*R*”) values of kmers (for *k* in the range 3-8 nt), where *R* is defined as the frequency of a kmer in protein-bound reads over its frequency in input reads (**Fig. 1A**, top right). In most cases, *R* values of top kmers exhibited a unimodal profile with increasing protein concentration consistent with increased signal above noise at moderate versus low RBP concentration, and increased binding of lower-affinity motifs at higher versus moderate concentrations (Lambert et al., 2014). A mean Pearson correlation across 5mers of 0.96 was observed among experiments performed on the same RBP at different concentrations, indicating high reproducibility (**Fig. 1D**). A comparison of previously reported binding specificities for 31 factors assayed by (Ray et al., 2013) using an independent array-based assay revealed high correlation with our dataset (**Fig. S1C,D,** mean Pearson correlation = 0.72), with only four proteins showing correlations below 0.5.

### Overlapping specificities of RNA binding proteins

In order to visualize and compare the primary sequence specificities of the assayed RBPs, we derived sequence motif logos for each RBP by aligning enriched 5mers (Z score ≥ 3, weighted by enrichment above input, using an iterative procedure that avoids overlap issues, **Fig. 1A** top right, **Methods**). For more than half of the RBPs (41/78), the enriched 5mers produced multiple distinct sequence logos, indicating affinity to multiple distinct motifs that may reflect different binding modes or binding by different RBDs (motif 5mers are listed in **Table S3**). Clustering of all RBPs by the similarity of their primary sequence logos revealed two clear trends: 1) the expected tight grouping of closely related paralogs (e.g., PCBP1/2/4, RBFOX2/3), (Conway et al., 2016; Smith et al., 2013); and 2) the more unexpected clustering of some completely unrelated proteins, often containing distinct classes of RBDs (**Fig. 2A**). Overall, 15 clusters of RBPs with highly similar primary motifs emerged (nine with three or more members) using a branch length cutoff chosen as described (**Methods**), while 18 RBPs with motifs more distinct from other profiled RBPs remained unclustered. Notably, eight of the 15 clusters contained two or more proteins with completely distinct types of RBDs (e.g., cluster 1 contained RRM-, KH- and ZNF-containing proteins as well as factors with multiple RBD types).

**Figure 2.**
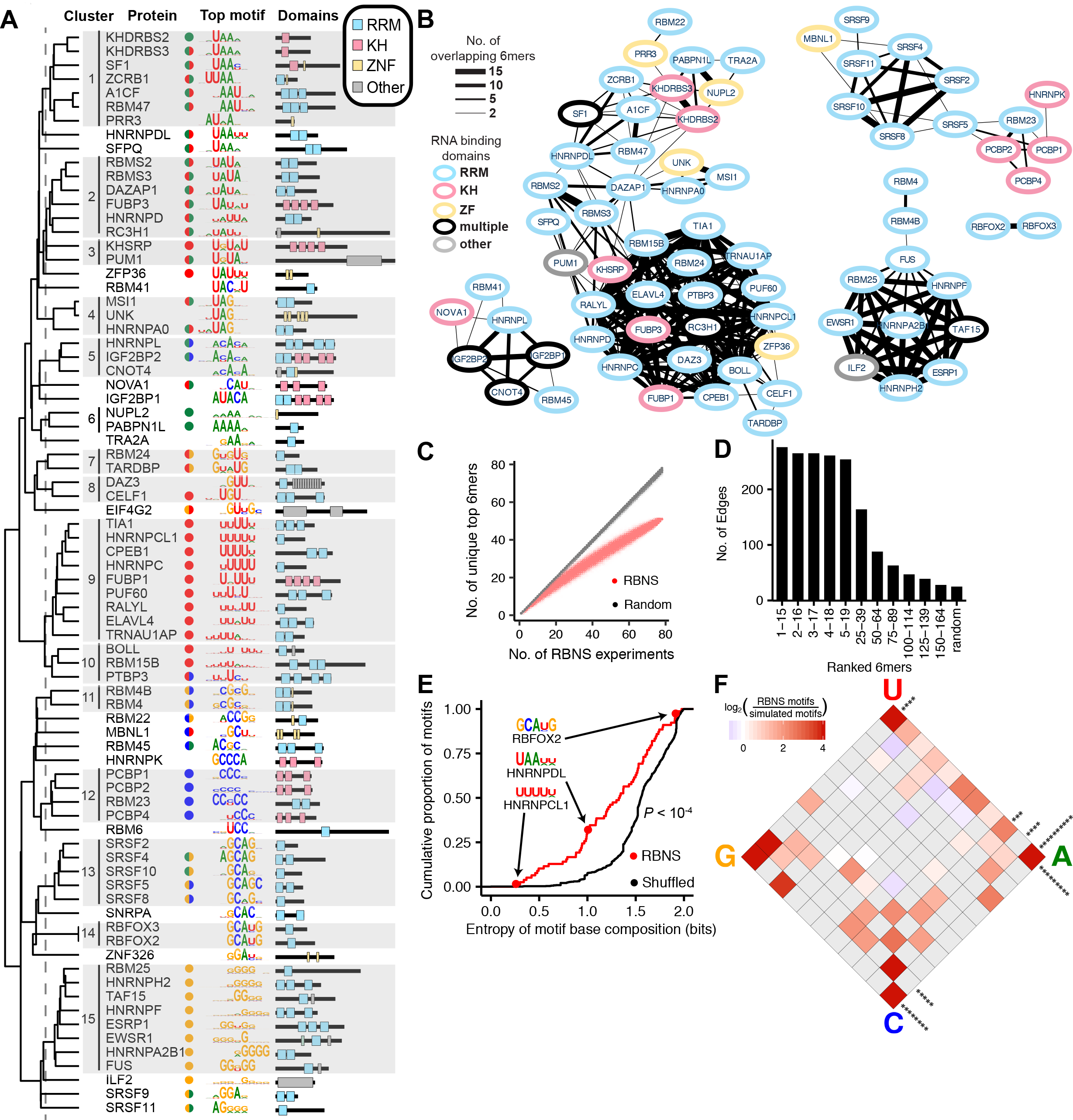
RBPs bind to a small subset of the sequence space, characterized by low-entropy motifs. **A.** From left to right: Dendrogram of hierarchical clustering of RBPs by sequence logo similarity and 15 demarcated clusters by branch length cutoff (dashed line); protein name; colored circles representing nucleotide content of RBP motif (one circle if motif is >66% one base, two half-circles if motif is >33% two bases); top motif logo for each protein; protein RBD. Each logo represented an average of seven 5mers. **B.** Network map connecting RBPs with overlapping specificities (sharing at least two of the top 15 RBNS 6mers). Line thickness increases with number of overlapping 6mers. Node outline displays RBD type of each protein. **C.** Number of unique top 6mers among subsamplings of the 78 RBNS experiments versus randomly selected 6mers. **D.** Edge count between nodes for network maps as shown in **B**, drawn using groups of 15 6mers with decreasing ranks. **E.** Entropy of nucleotide composition of RBNS motifs vs. simulated motifs (**Methods**). *P*-value determined by Wilcoxon rank-sum test. **F.** Enrichment of RbNs motifs over simulated motifs among partitions of a 2D simplex of the motif nucleotide composition (**Methods**). Significance along margins was determined by bootstrap Z-score (number of asterisks = Z-score).

To more rigorously assess the relatedness of RBP binding affinities, we generated a network map with edges connecting RBPs with significantly overlapping sets of top 6mers (at least two of the top fifteen 6mers, *P* = 0.001 hypergeometric test, **Fig. 2B**). While RBFOX2 and RBFOX3 were connected only to each other, other proteins were members of larger highly connected groups and the network overall was much more connected than expected (*P* < 10^−5^ relative to null distribution, **Methods**). Indeed, there were 27 overlaps among the #1 top 6mers of the 78 RBPs (collapsing to just 51 unique 6mers, **Fig. 2C**, red), compared to ~1 overlap expected by chance (**Fig. 2C**, black). A large excess of overlaps also occurred when considering the top fifteen 6mers for each RBP. Furthermore, the large excess of overlaps remained when eliminating clear paralogs and any proteins with at least 40% amino acid identity in an RBD to any other analyzed RBP (**Fig. S2A,B**). To explore the network’s connectivity further, we regenerated network maps with sets of 6mers of progressively decreasing affinities (e.g., 6mers ranked 2-16, 13-17, etc. for each RBP). A monotonic decrease in edges (overlaps of two or more) was observed with decreasing affinity categories, indicating that the connectivity of this RBP map is highest for 6mers with highest relative affinity (**Fig. 2D**). Together, these observations indicate that RBPs recognize a relatively small, particular subset of available sequence space. The pattern of clustering and overlap of motifs observed in Figures 2A-C, including many clusters of RBPs with distinct RBD types, implies that unrelated RBPs have evolved to bind similar RNA sequence motifs many times.

### RBPs preferentially bind low-complexity motifs

We noted that most of the RBP motifs identified were composed of just one or two distinct bases (**Fig. 2A**). To assess motif composition objectively, we measured the Shannon entropy of the nucleotide composition of each sequence logo, a scale which ranges from 0 bits (if motif composed 100% of one base) to 2 bits (25% each base). The entropies of actual RBP motifs were substantially lower than simulated motifs made from sampling columns from across RBPs (*P* < 10^−4^, Wilcoxon rank-sum test), indicating that RBP motifs are biased toward lower compositional complexity (**Fig. 2E**). This trend applied generally to all compositions with low complexity, as mapping RBP motif compositions onto a 2-dimensional simplex revealed increased density at all four mononucleotide “corners”, as well as all 6 dinucleotide “margins” (A/C, A/U, and C/U most significant, **Fig. 2F**, **Fig. S2C**, all bootstrap *P* < 0.05).

### RNA maps from RBNS and knockdown RNA-seq data

Many human RBPs are involved in pre-mRNA splicing. Therefore, it was not terribly surprising that ~35% of the 596 “RBNS 6mers” (those in the top 15 for any RBP) matched 6mer splicing elements identified previously in cell-based reporter screens (**Fig. 3A**, *P* = 1.7 × 10^−4^, hypergeometric test) (Ke et al., 2011; Rosenberg et al., 2015; Wang et al., 2012; 2013). In addition, the overlapping 6mers had stronger regulatory scores than non-RBNS 6mers (**Fig. 3B** left, *P* = 6×10^−36^, Wilcoxon rank-sum test), and higher RBNS enrichment (reflecting stronger binding) was associated with increased regulatory activity (**Fig. 3B** right for binned comparisons, overall Spearman ρ = 0.08, *P* < 10^−12^).

**Figure 3.**
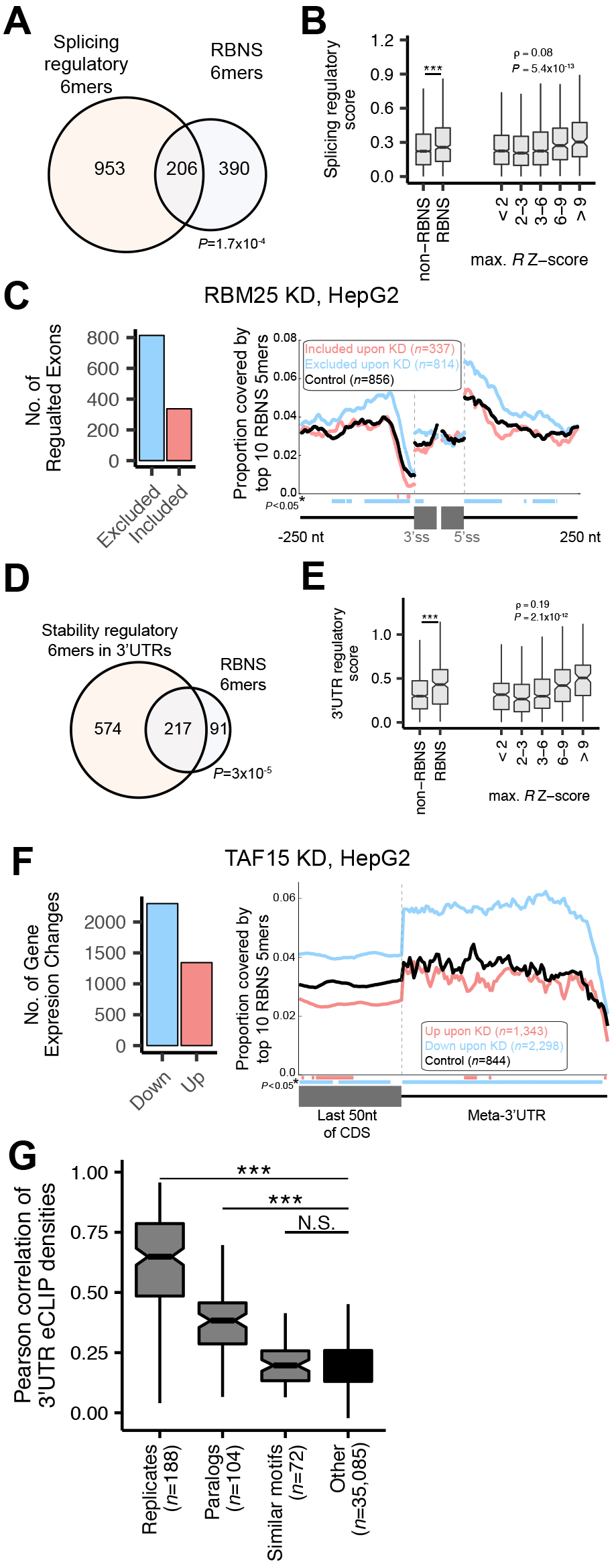
RBNS-derived motifs are associated with regulation of mRNA splicing and stability *in vivo*. **A.** Overlap of RBNS 6mers and 6mers with splicing regulatory activity (*P*-value determined by hypergeometric test). **B.** Comparison of splicing regulatory scores of, left: RBNS 6mers (top 15 of any RBNS, “RBNS”) and all other 6mers (“non-RBNS”); right: all 6mers binned by their maximum *R* value Z-score across all RBNS experiments (*P*-values determined by Wilcoxon rank-sum test). Overall Spearman correlation between *R* Z-score and splicing regulatory score was 0.08 (*P* < 10^−12^). **C.** Left: Number of alternative exons regulated by RBM25 as determined by RNA-seq after RBM25 knockdown in HepG2 cells. Right: Proportion of events covered by RBNS 5mers in exonic and flanking intronic regions near alternative exons excluded upon RBM25 KD (red), included by RBM25 KD (blue), and a control set of exons (black). Positions of significant difference from control exons upon KD determined by Wilcoxon rank-sum test and marked below the x-axis. **D.** Overlap of RBNS 6mers and 6mers with 3’ UTR regulatory activity (*P*-value determined by hypergeometric test among 1303 6mers with sufficient coverage for representation). **E.** Comparison of 3’ UTR regulatory scores of, left: RBNS 6mers (top 15 of any RBNS, “RBNS”) and all other 6mers (“non-RBNS”), *P*-value determined by Wilcoxon rank-sum test; right: all 6mers binned by their maximum *R* value Z-score across all RBNS experiments. Overall Spearman correlation between *R* Z-score and 3’ UTR regulatory score was 0.19 (*P* < 10^−11^). **F.** Left: Number of gene expression changes after knockdown of TAF15 in HepG2 cells. Right: Frequency of TAF15 RBNS 5mers along a meta-3’ UTR of genes whose expression is decreased (blue), increased (red) or unchanged (black) by TAF15 knockdown. Positions of significant difference from control genes upon KD determined by Wilcoxon rank-sum test and marked below the x-axis. **G.** Pearson correlations of eCLIP densities across 100nt windows of 3’ UTRs for all pairs of eCLIP experiments. Pairs of experiments are grouped by category, with all pairs not belonging to “Replicates”, “Paralogs”, or “Similar motifs” (sharing two of top 5 5mers) placed in “Other”. *P*-value determined by Wilcoxon rank-sum test, \*\*\**P*<5×10^−4^, N.S.=*P*>0.05.

“RNA maps” for splicing factors have traditionally been built using *in vivo* binding data from CLIP-seq combined with genome wide assays of splicing changes in response to RBP perturbation (Witten and Ule, 2011). To ask whether *in vitro* data could be used in place of CLIP data to derive such maps of inferred splicing activity, we integrated RBNS data with RNA-seq data from human K562 and HepG2 cells depleted of specific RBPs by shRNA (Van Nostrand et al., 2017). For example, depletion of RBM25 resulted predominantly in exclusion of cassette exons (**Fig. 3C**, left), and the G-rich motifs bound by this factor *in vitro* are enriched and conserved near splice sites (**Fig. S3A**). Constructing an RNA map for this factor based on its G-rich motif near significantly changing exons revealed an enrichment of top RBNS 5mers in introns flanking exons that were excluded upon RBM25 knockdown (KD) relative to control intronic sequences (**Fig. 3C**, right). Together, these data support that RBM25 functions as a splicing activator when it binds intronic motifs near alternative exons, consistent with previous reports showing that RBM25 controls splicing of specific alternative exons (Carlson et al., 2017; Gao et al., 2011; Zhou et al., 2008) and illustrate the potential of RBNS and RNAi/RNA-seq to provide informative splicing maps.

By performing this analysis on all 38 RBPs for which we had KD in at least one cell type with corresponding RBNS data, we observed that 27 of the 38 RBPs showed significant enrichment of their RBNS-derived 5mers in either activated or repressed exons or flanking introns, with enrichments often specific to intronic or exonic regions (**Fig. S3B**). Both splicing activation and splicing repression were inferred, involving for different RBPs both intronic and exonic regions (**Fig. S3C**). These RNA maps were consistent with previously reported roles of known splicing factors in many cases (e.g., splicing activation by DAZAP1 (Choudhury et al., 2014) and PUF60 (Page-McCaw et al., 1999) and repression by HNRNPC (Choi et al., 1986) and PTBP1 (Singh et al., 1995)), providing a fine-grained view of the position-specific nature of the RBP’s regulatory activity, bypassing the requirement for CLIP data. In some cases, RBPs not yet implicated in splicing regulation exhibited distinct motif enrichments in their RNA maps suggestive of function (e.g., ILF2 as a splicing activator). Of note, eight of the nine RBPs with G-rich motifs (FUS being the sole exception) exhibited splicing activator activity from at least one region (introns being most common), mirroring the G-rich *cis* regulatory sequences and candidate bound *trans* factors observed in unbiased screens for intronic splicing enhancers (Wang et al., 2012). Thus, this approach can provide a tool for understanding patterns of splicing regulatory activity that augments existing CLIP-based RNA maps (Van Nostrand et al., 2017).

### Protein-bound sequences are associated with *in vivo* regulation of mRNA levels

Besides splicing regulatory activity, we also observed significant overlap between RBNS 6mers and 6mers previously shown to modulate mRNA levels when inserted into reporter 3’ UTRs (Oikonomou et al., 2014) (**Fig. 3D**, *P* = 3×10^−5^, hypergeometric test). As observed for splicing regulation, 3’ UTR regulatory scores were higher for RBNS motifs (**Fig. 3E** left, *P* = 4×10^−10^, Wilcoxon rank-sum test), and regulatory scores increased for 6mers with higher RBNS enrichment (**Fig. 3E** right for binned comparisons, overall Spearman ρ = 0.19, *P* < 10^−11^).

Again using RNAi/RNA-seq data, we examined RBNS motif density in 3’ UTRs for genes significantly up- or down-regulated upon KD to generate 3’ UTR RNA maps. For example, TAF15 knockdown resulted in decreased levels of many mRNAs (**Fig. 3F**, left) and these mRNAs were enriched for TAF15 motifs in their 3’ UTRs relative to control genes with unchanged expression upon KD (**Fig. 3F**, right). These data suggest that TAF15 stabilizes mRNAs by binding to G-rich sequences in 3’ UTRs, mirroring expression changes observed upon TAF15 depletion in adult mouse brain and human neural progenitor cells (Kapeli et al., 2016). Just over half of the RBPs with corresponding KD data (20/38) had RNA expression maps that were consistent with a role in regulating mRNA levels (**Fig. S3D**), equally split between stabilizing and destabilizing activity. Interestingly, SRSF5 motifs were highly enriched in 3’ UTRs (and the end of the upstream ORF) of genes up-regulated upon KD (**Fig. S3E**). SRSF5 binding to 3’ UTRs has been observed in cell types in which this RBP undergoes nucleocytoplasmic shuttling (Botti et al., 2017), and its role in gene expression levels may be related to its role in linking alternative mRNA processing to nuclear export (Müller-McNicoll et al., 2016).

Sequence elements involved in post-transcriptional regulation are often more frequent and more evolutionarily conserved in transcript regions where they function (Lim et al., 2011; Xie et al., 2005). Considering the set of all 5mers that were bound by at least one factor, this set was more conserved than non-RBNS 5mers in intronic regions near splice sites (**Fig. S4A**, **Methods**). This effect was greater in introns near alternative exons and even stronger in introns flanking deeply conserved alternative exons (**Fig. S4A**, *P* < 0.05, Wilcoxon rank-sum test), consistent with the presence of many important regulators of alternative splicing among the assayed RBPs. Enrichment and conservation was also observed in other specific transcript regions (**Fig. S3A**) (Fairbrother et al., 2002; Goren et al., 2006; Voelker and Berglund, 2007). For example, the EIF4G2 RBNS motif was enriched and conserved only in the 5’ UTRs of genes, consistent with its known role in translation initiation, while the motif of the splicing and stability factor TIA1 was enriched and conserved in both introns and 3’ UTRs of transcripts, and many other cases of enrichment in transcript regions associated with known RBP functions were observed (**Fig. S3A**). Thus, for RBPs of unknown function the pattern of RBNS motif enrichment and conservation can also be used generate hypotheses about function.

### RBPs with similar motifs can target and regulate different events

As part of a larger analysis of ENCODE RBP data, we compared RBNS motifs to *in vivo* binding patterns assessed by eCLIP when such data were available (Van Nostrand et al., 2017). We observed strong agreement between eCLIP and RBNS motifs in most cases, with 17 of 28 proteins having significant overlap between RBNS 5mers and 5mers identified *de novo* as enriched in eCLIP peaks (**Fig. S4B**, adapted from (Van Nostrand et al., 2017)). Furthermore, RBNS-enriched 5mers were more enriched in eCLIP peaks identified in multiple eCLIP replicates and in peaks identified in multiple cell types, which likely represent sites of more robust binding (**Fig. S4C**). Together, these observations support that RBNS-identified motifs drive the RNA binding specificity of most RBPs.

In cells, RBPs appear to bind only a subset of cognate motifs in expressed transcripts (Taliaferro et al., 2016), and the extent to which RBPs with similar binding motifs bind the same targets *in vivo* is incompletely understood. Analyzing eCLIP data for 131 RBPs, we observed somewhat elevated correlation of binding locations between highly similar paralogs (**Fig. 3G**), with mean Pearson correlation of 0.37, relative to correlations of ~0.2 between randomly chosen proteins, but far below that observed between replicate eCLIP experiments (mean correlation = 0.67). However, despite the generally similar motif enrichments observed *in vivo* and *in vitro*, we observed surprisingly little correlation between binding locations of other pairs of RBPs that bound similar motifs *in vitro* (sharing at least two of their top five RBNS 5mers) (**Fig. 3G**), with a mean Pearson correlation of 0.20, not different from random pairs of RBPs. For example, while TIA1 and HNRNPC both have high affinity for U-tracts *in vitro* and *in vivo*, they bind distinct sites in many transcripts (example shown in **Fig. S4D**).

The low correlation between *in vivo* binding sites of RBPs with similar motifs could result from various factors, including: i) differences in subcellular localization of RBPs resulting in differential access to transcripts; ii) differential participation in complexes with other factors that alter RNA specificity (e.g. (Damianov et al., 2016)); iii) occlusion of sites by one RBP leading to the exclusion of other RBPs (Zong et al., 2014); iv) technical differences in efficiency of eCLIP capture of different regions by different RBPs; or v) subtler differences in binding specificities not well captured by conventional motif representations resulting in binding different subsets of motif instances. While all of these factors likely contribute to some extent, we focused here on exploring the fifth possibility, leveraging the depth and sensitivity of the RBNS data to explore binding determinants beyond canonical short RNA motifs.

### RNA structural preferences of RBPs

The RNA secondary structure of RNA motifs can impact RBP binding and regulation (Warf et al., 2009), modulate the efficacy of regulatory RNA sequences (Hiller et al., 2007), and improve *in vivo* RBP binding site prediction (Li et al., 2010). Since potential RBP binding sites in the transcriptome exist in a variety of structural conformations, we determined RNA secondary structure preferences for each RBP by computationally folding ~5 million input and compositionally-matched protein-bound reads for all 78 experiments at all concentrations (**Methods**). To assess the RNA secondary structure preferences of all RBPs assayed, we first calculated the base pairing probability (*P*_paired_) of occurrences of the top RBNS 6mer in pulldown libraries as well as positions flanking each motif occurrence (**Fig. 4A**). Individual factors displayed a range of structural preferences with most proteins disfavoring structure at most motif positions compared to flanking positions, with an average ~20% decrease in base pairing probability over the center of the motif relative to flanking sequences. Examining RBPs by motif similarity clusters as shown in Figure 2A, RBPs that bound AU-rich (clusters 1, 2) and A-rich (cluster 6) motifs disfavored structure most strongly, while factors binding U-rich motifs (clusters 9, 10) disfavored structure more moderately, and C-rich (cluster 12), G-rich (cluster 15), and CG-rich (cluster 11) motifs tended to be just as or more structured than their flanking positions. Most RBPs also preferred relatively unstructured occurrences of their motifs (pulldown *P*_paired_ < input *P*_paired_, **Fig. 4A**, right bar), with just six preferring structure when averaging over the entire motif, with the strongest preference observed for ZNF326.

**Figure 4.**
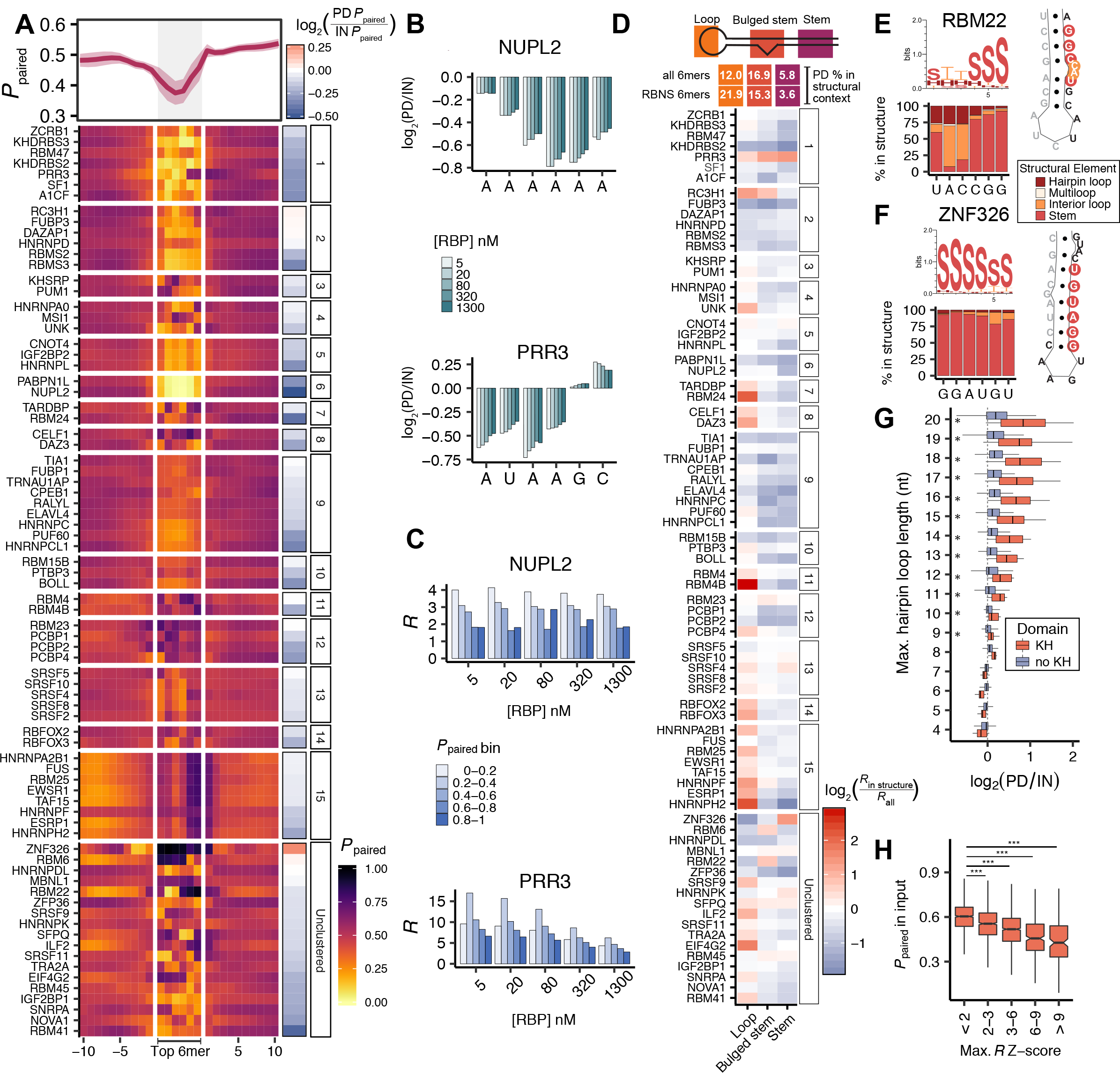
RNA secondary structural preferences of RBPs. **A.** Top: *P*_paired_ over each position averaged over the 78 RBPs; 95% confidence interval is shadowed. Bottom: Mean *P*_paired_ in the most enriched pulldown library over the top 6mer plus 10 flanking positions on each side; RBPs are grouped by motif clusters in **Fig. 2A**. Right: Mean change (log_2_) in pulldown vs. input *P*_paired_ averaged over the top 6mer. **B.** Mean change (log_2_) in *P*_paired_ over each position of the top 6mer at different concentrations of NUPL2 (top) and PRR3 (bottom) relative to the input library. **C.** Enrichment of the top 6mer of NUPL2 (top) and PRR3 (bottom) in 5 bins into which all 6mers were assigned based on their average *P*_paired_. **D.** Top: Three types of structural contexts considered and the percentage of all 6mers and RBNS 6mers (top 6mer for each of 78 RBPs) found in each context in pulldown reads. Bottom: Log-fold change of the top 6mer’s recalculated *R* among 6mers restricted to each structural context relative to the original *R*. **E, F:** Left: Percentage of each position of the top 6mer found in the four structural elements for RBM22 (**E**) and ZNF326 (**F**) in pulldown reads. Structure logo for top 6mer is shown above. Right: Representative MFE structures of the top 6mer pairing with the 5’ sequencing adapter (gray) for 6mers found at the most enriched positions within the random 20mer (RBM22, position 5; ZNF326, position 6). **G.** Enrichment of the percentage of pulldown vs. input reads containing hairpin loops of various lengths, separated by RBPs that contain (*n*=13) vs. do not contain (*n*=65) at least one KH domain. Lengths with significant differences determined by Wilcoxon rank-sum test (*P*<0.05). **H.** Average *P*_paired_ in input libraries for all 6mers binned by maximum *R* value Z-score across all RBNS experiments (\*\*\**P*<10^−22^ by Wilcoxon rank-sum test; Spearman correlation =-0.4, *P* < 10^−15^).

Many RBPs showed varying degrees of secondary structure preferences at different positions within the top 6mer (**Fig. 4A**). To explore this observation further, we calculated the mean base pair probability at each position within bound 6mers relative to that of the same 6mer in input libraries. For example, NUPL2, a protein that binds A_6_ motifs, strongly disfavored structure at all positions (**Fig. 4B**, top), whereas PRR3 had a preference for positions 5 and 6 of its top AUAAGC motif to be paired but positions 1-4 unpaired (**Fig. 4B**, bottom).

To assess the effect of secondary structure on enrichment, we recomputed *R* values for all 6mers considering only occurrences of the 6mer in five contexts ranging from unstructured (average *P*_paired_ < 0.2 over the 6 positions) to structured (average *P*_paired_ ≥ 0.8) (**Fig. 4C**, **Methods**). Consistent with the notion that PRR3 prefers a partially structured motif while NUPL2 does not, PRR3’s top 6mer was most enriched in a moderately structured context (*P*_paired_ 0.2-0.4, **Fig. 4C**, bottom) while NUPL2’s top 6mer was most enriched in the least structured context (*P*_paired_ 0-0.2, **Fig. 4C**, top). Furthermore, for PRR3 and NUPL2, the *R* values of the top 6mer were 3-and 4-fold higher, respectively, in the most enriched *P*_paired_ bin relative to the least enriched *P*_paired_ bin, underscoring the impact of RNA secondary structure on affinity.

As particular RNA structures are known to affect RBP binding in pre-mRNA splicing (Warf and Berglund, 2010), mRNA decay (Goodarzi et al., 2012), and mRNA localization (Rabani et al., 2008), we classified each RNA base as being part of a stem, hairpin loop, interior loop, or multiloop (Kerpedjiev et al., 2015). Averaging over all 78 proteins, we found that in pulldown sequences, top RBNS 6mers were present in hairpin loops about twice as often as all 6mers and in stems approximately half as often (**Fig. 4D**, **top**). Correspondingly, the top 6mers of many individual RBPs were more enriched in a loop context (**Fig. 4D**, bottom), including GU-binders (clusters 7, 8), CG-binders (cluster 11), RBFOX2/3 (cluster 14), and almost all G-rich binders (cluster 15). Fewer RBP motifs were preferentially enriched in a bulged stem (9 RBPs) or stem (8 RBPs) context, with generally more modest enrichments than seen for hairpin loops (all enrichments reported in **Table S4**). Among the strongest bulged stem and stem-preferring RBPs were the core spliceosomal protein RBM22 (**Fig. 4E**) – which binds catalytic RNA structural elements in the spliceosome and makes direct contacts with the U6 snRNA Internal Stem Loop and intron lariat (Rasche et al., 2012; Zhang et al., 2017) – and the zinc-finger protein ZNF326 (**Fig. 4F**). Unlike most other RBPs, the motifs for these two proteins showed uneven distributions along sequence reads, with increased frequency at the 5’ end of the random sequence, and were commonly predicted to basepair with the 5’ sequencing adapter (**Fig. S5A**). Thus, it seems probable that RBM22 and ZNF326 primarily recognize RNA secondary structures, with specific motifs enriched because they can form these structures by basepairing with the sequencing adapters used, but not conferring sequence specificity in other contexts.

Analyzing the structural preferences of RBPs containing different RBD types, the most striking pattern was an enrichment of hairpin loops in sequences bound by RBPs containing KH domains (**Fig. 4G**). This enrichment was observed for 10/13 KH-containing RBPs, including all of those assayed except the three members of the FUBP family that bind U-rich sequences. Given that most (7/10) of these RBPs contain more than one KH domain, it is possible that relatively large hairpin loops allow binding of multiple KH domains to the RNA as has been observed in a crystal structure of NOVA1 (Teplova et al., 2011) and in SELEX analysis of PCBP2 (Thisted et al., 2001).

For RBPs for which we had corresponding eCLIP data (*n* = 27), we observed a high correlation between RNA secondary structure preferences *in vitro* and *in vivo* (**Fig. S5B**). While most RBPs disfavored structure in both assays, RBPs that were found to prefer structured or partially structured motifs in RBNS, such as RBM22 and SFPQ, showed this same preference *in vivo*.

One puzzle raised by our observation above that human RBPs preferentially bind a small subset of motifs is why this particular subset of motifs has been chosen. Analyzing RNA folding in the input RNA pool consisting of essentially random RNAs, we noted that top RBNS motifs tend to be less structured even in input reads than are other 6mers. Furthermore, 6mers with higher RBNS enrichment were even less structured (**Fig. 4H** for binned comparisons, overall Spearman correlation = −0.39, *P* < 10^−13^). Given that most RBPs prefer binding to unstructured motif instances, as observed previously and above (e.g., **Fig. 4A**), this observation suggests that many RBPs have evolved specificity for motifs that are intrinsically less structured, perhaps to enable binding to a larger fraction of motif occurrences in the transcriptome.

### RNA binding proteins interact with short spaced motifs

Although structural studies have described a variety of ways that RBDs engage RNA, it is generally thought that a single RBD (e.g., an RRM or KH domain) makes contacts with 3-5 contiguous RNA bases (Auweter et al., 2006). Of the factors in this study, more than half contained multiple RBDs (**Fig. 1B**) or multiple types of RBDs, raising the possibility that these RBPs can interact with pairs of small motifs spaced one or more bases apart, hereafter referred to as ‘bipartite motifs’. For example, NMR structures of MSI1’s individual RRMs revealed they bound GUAG and UAG respectively, leading to a structural model where both RRMs bind the sequence UAGN(0-50)GUAG together (Iwaoka et al., 2017).

RBNS is well suited for the unbiased identification of bipartite motifs due to the complex sequence space and depth of sequencing, as longer spaced Aimers are well represented. We computed enrichments for motifs composed of two cores of length 3 bases separated by spacers of length 0-10 (**Fig. 1A**, **Methods**) (where spacing 0 represents a traditional contiguous 6mer motif). No preference for spaced motifs relative to linear motifs was detected in control experiments using this method (**Fig. S6A**), indicating that bipartite and linear motifs of equal information content but different lengths can be directly compared.

This analysis yielded many RBPs enriched for bipartite motifs with varying spacing preferences. For example, we found that the dual RRM-containing DAZAP1 protein preferred AUA followed by another AUA-containing core spaced by 1-3 nucleotides, with no particular preference for any of the four bases in the spacer (**Fig. 5A**). The three RRM protein RBM45 bound two AC cores, separated by a spacer of 1-3 nucleotides, with a bias against Gs in the spacer (**Fig. 5B**). Analysis of all 78 factors revealed that almost one third of RBPs bound bipartite motifs with similar or greater affinity than linear 6mers, with 17 RBPs showing a significant preference for a spaced over linear motif at a 5% FDR (**Fig. 5C**, **Methods**).

**Figure 5.**
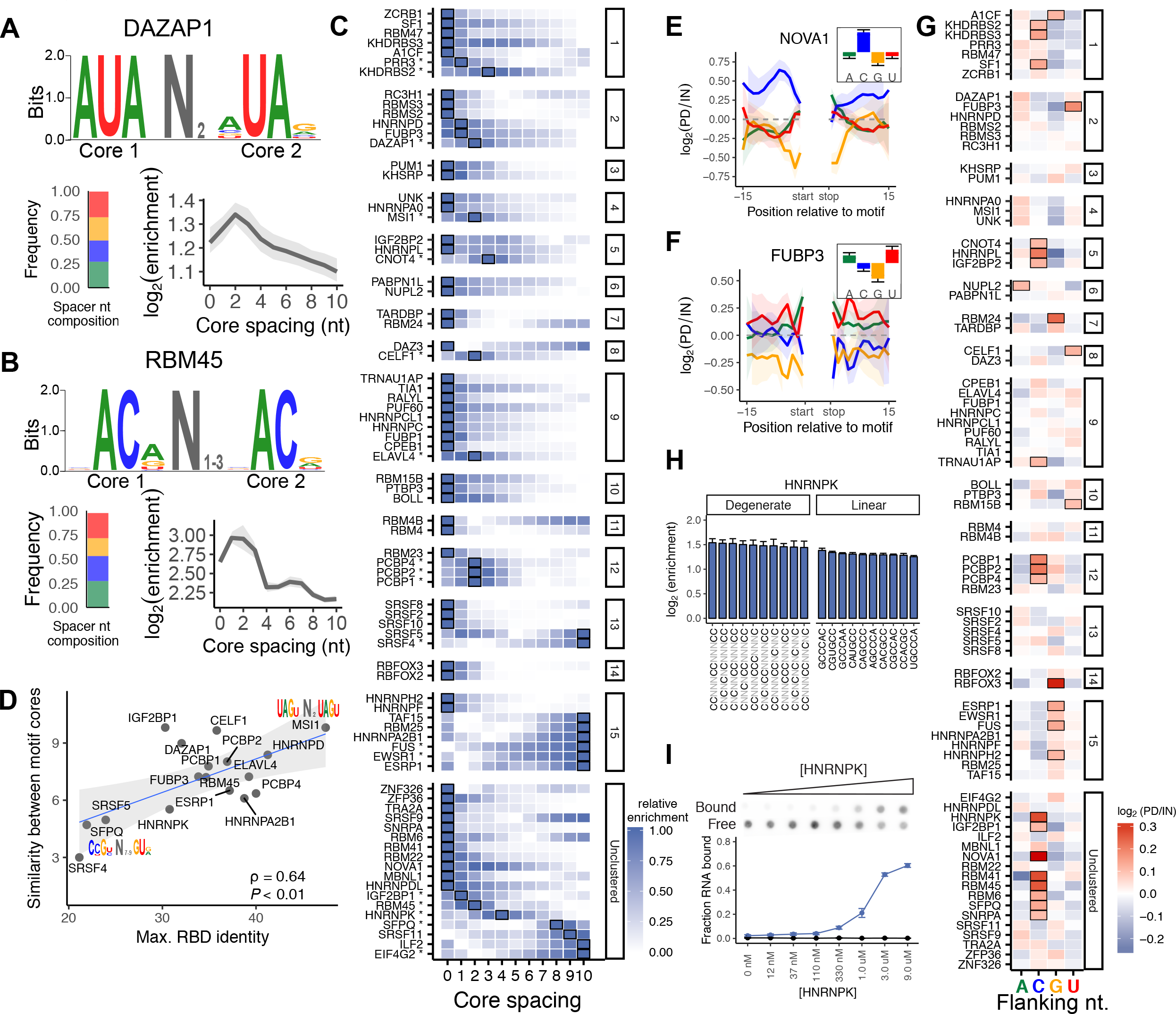
Many RBPs bind bipartite motifs or prefer flanking nucleotide contexts. **A, B.** Top: Sequence logos of bipartite motifs for DAZAP1 (**A**) and RBM45 (**B**). Bottom: Nucleotide composition of the spacer between both motif cores (left) and enrichment as a function of the spacing between cores (right). **C.** Core spacing preferences of all RBPs. Each row indicates the relative enrichment as a function of the spacing between cores for a given RBP (i.e., enrichments normalized to the maximum in each row). A box indicates the spacing with maximal enrichment for that RBP, and * to the right of the RBP name signifies the non-zero spacing is significantly preferred over the best linear 6mer. RBPs are grouped by motif clusters in **Fig. 2**. **D.** Pearson correlation between the maximum similarity of RBDs of the same type within an RBP and the similarity between the core motifs of the best bipartite motif. Only RBPs with a significant preference for spacing greater than 0 in **C** and those with at least two RBDs of the same type were used. **E. F.** Flanking nucleotide context preferences surrounding the top five NOVA1 (**E**) and FUBP3 (**F**) 5mers. Reads with no secondary motifs were centered around the top 5mer and the enrichment for each nucleotide in protein-bound reads relative to input reads was calculated (**Methods**). Inset: mean enrichments across all positions flanking the motif for each of the four nucleotides. **G.** Flanking nucleotide context preferences of all RBPs. Each row displays the enrichment or depletion for each nucleotide surrounding the RBP’s top five 5mers. Boxes indicate significant enrichment or depletion (|log_2_(enrichment)|>0.1 and *P*<0.001, **Methods**). **H.** Enrichments of HNRNPK’s top 10 linear 6mers (right) and top 10 degenerate sequences of length 12 with 6 Ns (left). **I.** Filter assay validation of HNRNPK binding to the oligo UUU(CCUCUCUUUUCC)UUU (blue) and the oligo U_12_ (black) as a negative control (**Methods**). Dot-blot of filter assay shown on top with fraction of RNA bound quantified below.

Many RBPs with multiple RRMs that bound bipartite motifs preferred a short spacer of 1-4 nucleotides, consistent with the RNA spacer lengths of solved structures of tandem RRMs with a single RNA, such as CELF1 and Sex-lethal (Afroz et al., 2015; Kanaar et al., 1995; Teplova et al., 2010). As expected, preference for spacing was higher for factors that contained more than one known RBD (**Fig. S6B**, *P* < 0.023, t-test), although several exceptions were observed. For example, KHDRBS2 showed the greatest preference for a bipartite motif even though it only has just one KH domain (**Fig. S6B**). However, KHDRBS2 includes an N-terminal QUA1 (Quaking-1) domain, a domain which has been shown to promote homodimerization of some STAR (Signal Transduction and Activation of RNA) family proteins (Beuck et al., 2012; Meyer et al., 2010), suggesting that KHDRBS2 may bind bipartite motifs as a homodimer.

We also observed several proteins whose affinity for RNA increased with longer spacings, exhibiting greatest enrichment at the maximal allowed spacing of ten bases (e.g., TAF15 and ESRP1 in cluster 9 of **Fig. 5C**, **Fig. S6C**). One potential explanation may be RNA-dependent multimerization. For example, FUS, a factor that displayed increasing enrichments as a function of spacing, has a C-terminal RG-rich domain that has been shown to promote cooperative binding to RNA (Schwartz et al., 2013). Notably, EWSR1, a FUS paralog that is a member of the FET (FUS, EWSR1, TAF15) family and has a similar domain composition (Schwartz et al., 2015), displayed the same preference for increased spacing, suggesting it may also multimerize through low-complexity domains.

We noted that the two RNA cores bound by MSI1 were nearly identical, and MSI1’s two RRMs were highly similar at the amino acid level (~47% identity), while SFPQ favored a motif consisting of two very different RNA cores, and it’s RBDs were much less similar (~22% identical). Expanding this observation to all RBPs that preferred spaced motifs and contained at least two RRM, KH, or ZF domains, we observed a fairly strong positive correlation between the percent identity of sibling RBDs within a protein and the similarity of the bipartite motif RNA cores (**Fig. 5D**, Pearson correlation = 0.64, *P* < 0.01). These observations support the model that most binding of bipartite motifs in the set of RBPs analyzed involves engagement of RNA by more than one RBD.

### RNA sequence context enhances RBP binding

It has previously been observed that binding of certain transcription factors may be enhanced by particular nucleotide composition(s) adjacent to a high-affinity motif (Jolma et al., 2013) and that such flanking nucleotide biases are also seen around motifs within ChIP-seq peaks (Wei et al., 2010). We hypothesized that adjacent nucleotide context could play a role in modulating RBP specificity by potentially altering local RNA secondary structure or creating additional interactions with the RBP. One such example is Argonaute-2, which preferentially binds miRNA target sites in an AU-rich context, a feature often used to predict miRNA targeting efficacy (Agarwal et al., 2015; Grimson et al., 2007; Nielsen et al., 2007). For each RBP, we computed the enrichment of each nucleotide at all positions in reads surrounding a high-affinity motif, using only those reads containing one of the top five 5mers and no secondary motifs (**Methods**). We found 28 proteins with a significant preference for a particular flanking nucleotide context (mean log_2_(enrichment) > 0.1, *P* < 10^−3^ by Wilcoxon rank-sum test, **Methods**). For example, NOVA1 preferred to bind its motif in a C-rich context (**Fig. 5E**) while FUBP3 preferred to bind its motif in a U-rich context (**Fig. 5F**). We noted an enrichment for RBPs with KH domains within this set (*P* < 10^−3^, Fisher’s exact test), as seen for factors that prefer binding to RNAs within hairpin loops (**Fig. 4H**). While particular flanking nucleotide compositions may be correlated with the presence of large hairpin loops, we observed a majority of these flanking nucleotide context preferences even after controlling for the secondary structure context of the motif, suggesting that nucleotide context effects and secondary structure both contribute to binding (**Fig. S6D**). In most cases, this nucleotide preference was dependent on the presence of a motif in the read, suggesting that flanking sequence promotes or stabilizes RBP binding to a primary motif. However, some RBPs showed these same nucleotide preferences in the absence of a high-affinity motif (e.g., FUS, IGF2BP1, **Fig S6E**), suggesting that these factors have some affinity for degenerate sequences with biased nucleotide content.

To formalize these observations and test cases in which biased sequence composition might better describe an RBP’s specificity than a linear motif, we calculated enrichments for degenerate patterns with biased nucleotide composition. For example, HNRNPK, one of the RBPs that showed a preference for C bases in the absence of a high-affinity *k*mer, had greater enrichment for the degenerate pattern CNCNCNCNNNCC (enrichment=2.9) than the corresponding contiguous 6mer CCCCCC (enrichment=1.11) with identical information content of 12 bits. In fact, HNRNPK showed greater or equal enrichments for various permutations of the above degenerate pattern (6 Cs with 6 interspersed Ns) relative to the most highly enriched linear 6mers (**Fig. 5H**). We generalized this observation by calculating enrichments for all degenerate patterns composed of 6 fixed bases and 6 Ns for each RBP (**Methods**) and found that HNRNPK had higher enrichment for many C-rich degenerate patterns than for its best linear *k*mers of equal information content, whereas most other RBPs such as RBFOX2 strongly preferred specific linear sequences over degenerate patterns (**Fig. S6F**).

We selected a highly enriched degenerate pattern that was predicted to bind HNRNPK (CCNCNCNNNNCC). Substituting U at the N positions in order to avoid creating RNA secondary structure or other potential motifs, we validated that HNRNPK specifically bound to RNAs containing this pattern using a filter binding assay (**Fig. 5I**). These degenerate patterns were also enriched more than two-fold relative to linear 6mers in HNRNPK eCLIP peaks, supporting binding *in vivo* (**Fig. S6G**). In all, we identified 17 RBPs whose binding was well described by degenerate patterns, 14 of which bound spaced motifs in **Fig. 5C** and were often enriched for patterns similar to the identified bipartite motifs (e.g., CELF1, **Fig. S6H**). However, some RBPs showed enrichment for patterns with no more than 2 contiguous specified bases (e.g. FUBP1, **Fig. S6H**). These patterns may therefore represent degenerate bi- or tri-partite motifs, where binding of multiple RBDs, each specifically contacting just 1 or 2 RNA bases, results in a flexible binding pattern allowing the RBP to recognize an unusually broad range of RNA targets. Bipartite motifs and nucleotide context preferences for all 78 RBPs are listed in **Table S5**.

### Towards a more complete characterization of RBP specificities

Our observation of common preferences for bipartite motifs, flanking nucleotide context and secondary structural features led us to hypothesize that RBPs that favor similar motifs may often diverge in their preferences for these other features in order to impact different subsets of targets. For example, PCBP1 and RBM23 (cluster 12) both bind C-rich sequences even though they have distinct RBD composition (3 KH domains versus 2 RRMs). Our analysis indicated that PCBP1 disfavors structure over its motif, is capable of binding the bipartite motif CCCNNCCC, and is enriched for flanking C bases. In contrast, RBM23 has no structural preference over its motif and favors a contiguous C-rich motif with no flanking nucleotide context preference. Thus, PCBP1 and RBM23 are likely to bind distinct populations of C-rich sites in transcripts.

In order to systematically compare the contributions of these features, we calculated “feature-specific” *R* values by calculating the top motif’s *R* value as a function of: i) *P*_paired_ of the 6mer; ii) the average base pairing probability of the sequence flanking the 6mer (*P*_flank_); or iii) the nucleotide frequencies surrounding the 6mer (**Methods**). Since bipartite motif enrichment does not require presence of a linear motif, feature-specific *R* values for bipartite motifs were calculated based on the pattern of preference for spacings between the split motif cores (analogous to **Fig. 5C**, **Methods**). We then measured an information-theoretic distance between feature-specific *R* value profiles for pairs of RBPs within the same motif cluster (**Fig. S7A**, **Fig. 6A**). Intra-cluster pairwise distances were significantly higher than distances calculated between replicate RBNS experiments at different RBP concentrations for *P*_paired_, context and bipartite motifs. This observation suggests that RBPs with similar primary motifs are often differentially affected by contextual features (**Fig. 6A**).

**Figure 6.**
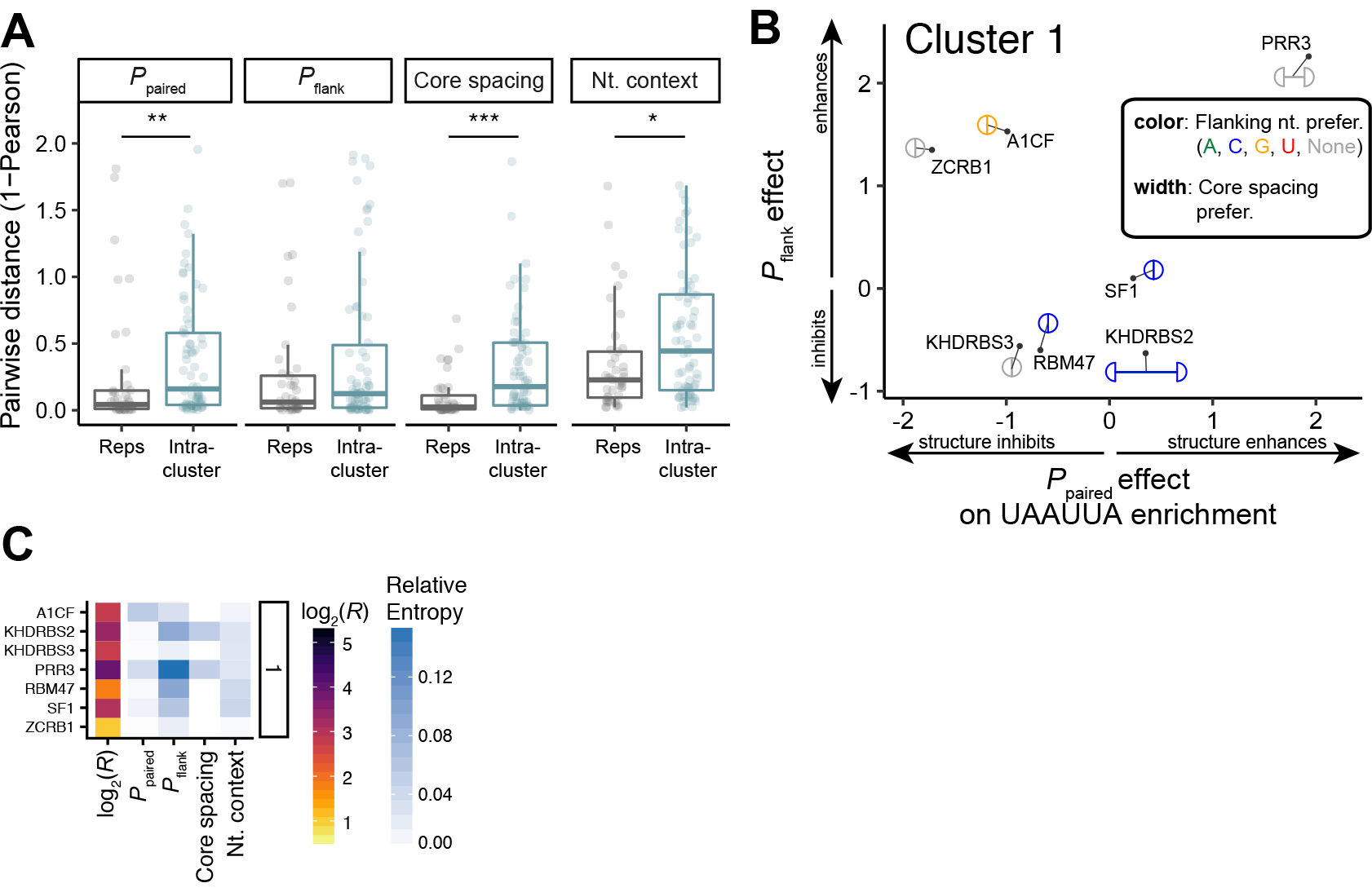
RBPs that bind similar motifs diverge in sequence context preferences. **A.** Pairwise distances (1-Pearson) of feature-specific *R* values for pairs of RBPs within a motif cluster (“Intra-cluster”) compared to distances between controls (“Reps”), where controls are replicate assays of the same RBP done on different days and at different protein concentrations. *P*-values determined by Wilcoxon rank-sum test (*< 0.05, **0.005, ***0.0005). **B.** Overall dispersal of specificities between cluster 1 RBPs for UAAUUA. x- and y- axes represent the degree to which secondary structure over (x) and flanking (y) the motif affect UAAUUA *R* values. Coloring of circle denotes whether the protein displayed a significant enrichment for flanking nucleotide context, with gray denoting none. Split circles indicate whether the protein had significant preference for a bipartite motif over a linear motif with the distance between the half-circles reflecting the preferred spacing of the cores. **C.** For cluster 1 RBPs: Left: The maximum *R* value (the greater of the best linear or bipartite motif) for each RBP. Right: The relative entropy of the distribution of pulldown vs. input top 6mer occurrences in five context bins for each of the context features (*P*_paired_, *P*_flank_, core spacing, and flanking nucleotide context).

To visualize protein preferences for each of the four features among RBPs within the same cluster, we placed RBP markers (consisting of paired semicircles) on a coordinate system according to their structural preferences *P*_paired_ and *P*_flank_ and then colored each RBP marker based on its flanking nucleotide preferences, and separated the semicircles based on the bipartite spacer preferences (**Methods**). This visualization of RBPs within the AU-rich cluster 1 revealed that no two RBPs are superimposed in this multidimensional affinity space (**Fig. 6B**), and divergences were observed within most other clusters as well (**Fig. S7B**). Overall, we found that 9/15 clusters diverged significantly in at least one feature, and 5/15 diverged in more than one feature with the most common significant feature being bipartite motif spacing (**Table S6**).

Using a relative entropy measure (**Methods**) we quantified the amount of information conveyed about a motif’s context by the knowledge that it is bound by a given RBP, allowing comparison of all features on the same scale within and between proteins. As shown in **Fig. S7C**, relative entropy values for the structure of the motif, the structure flanking the motif, the core spacing of bipartite motifs, and the nucleotide context for each RBP varied from zero for features that did not differ between pulldown and input motifs to ~0.6 bits for the *P*_paired_ context of ZNF326, which strongly prefers binding to a stem (**Fig. 4F**). Considering the AU-binding RBPs of cluster 1 as an example, the context feature with greatest effect on PRR3 binding was structure flanking the UAAUUA motif (**Fig. 6C**), with motifs in pulldown reads having much less flanking secondary structure than input occurrences (**Fig. S7D**). This and other context effects impact the overall specificity of this and other RBPs in the cluster significantly (**Fig. 6C**, left; **Fig. S7C**, top). In summary, we find that the specificity of most RNA binding proteins is conferred not only through canonical primary sequence elements but also by a variety of contextual properties of the local RNA sequence environment.

## DISCUSSION

### Towards a parts list of RNA regulatory sequence elements and *trans*-acting proteins

Over the past decades a large body of work has aimed to catalog functional RNA elements and their interacting proteins to gain a more complete and mechanistic understanding of RNA processing in cells. For example, analysis of *k*mer frequency and evolutionary conservation in specific regions of the genome has led to the appreciation that many *cis* sequences are associated with specific types of regulation (Fairbrother et al., 2002; Ke et al., 2011), even if the underlying *trans*-acting RBPs that bind them are not always known. Here, we characterized 78 human RBPs using a high-throughput version of RNA Bind-N-Seq, a one-step *in vitro* binding assay that assesses the spectrum of RBP binding specificities along with features that influence binding such as secondary structure, flanking nucleotide context and bipartite motifs.

### RBPs recognize a small subset of the available sequence space

Considering a diverse set of 78 factors, we find that many RBPs bind a relatively small, defined subset of primary RNA sequence space rich in low-complexity motifs primarily composed of one or two bases. This trend was seen independently for RRM and KH domains that do not share common ancestry, suggesting the existence of evolutionary pressures on RBPs to recognize these kinds of motifs. These findings are consistent with previous studies implicating AU-, U-, and G-rich sequences as functional elements that regulate stability and splicing (Fu and Ares, 2014; Wu and Brewer, 2012). Previous studies have shown certain mono- and dinucleotide rich sequences occur in clusters to mediate their effects on RNA processing (Barreau et al., 2005; Cereda et al., 2014). These sequences exhibit greater robustness to mutation than more complex sequence motifs. For example, RBPs in cluster 9 bind more than 50% of 5mers that differ by one from their top U_5_ motif whereas unclustered RBPs bind only 15-20% of 5mers that differ by one from their highest-affinity 5mer. Furthermore, these sequences also have lower propensity for strong secondary structures both in random sequence (**Fig. 4H**) and in the transcriptome, which may contribute to their choice as common RBP targets. Mechanisms involving sliding of RBPs along RNA may also favor low complexity motifs. For example, HNRNPC was shown to bind runs of uridines with a potential to slide along different registers within the RNA (Cienikova et al., 2014). Such a sliding model would be most feasible for mono-or dinucleotide repeat motifs because more complex motifs would likely require the RBDs to completely dissociate from the RNA to find another motif occurrence.

A similar analysis of DNA-binding proteins and transcription factors (not shown) revealed that they do not show the same inherent propensity to target low complexity sequences, perhaps due to differences in the size of the search space and differences in DNA and RNA structure and the biochemical mechanism of binding (Jolma et al., 2013). While the overlapping specificity across these RBPs is high we find that 18 of the 78 profiled RBPs have motifs dissimilar from that of any other RBP assayed (e.g., SNRPA), suggesting presence of a subset of RBPs that may have evolved to recognize a more specialized set of RNA targets. Indeed, these 18 RBPs tended to have broader expression profiles than RBPs within motif clusters (**Fig. S7E**), while RBPs with motifs shared by other RBPs are more likely to be tissue-specific. Thus, tissue specificity may reduce the number of RBPs targeting similar motifs present in any particular cell type or state. Even so, coexpression of many members of a cluster appears widespread, and this is likely to contribute in important ways to the function of post-transcriptional gene regulatory networks.

### RBP binding specificities harbor hidden complexity

Closer analysis of the complexity of RBNS data revealed that linear sequence is often insufficient to fully capture RBP binding specificities, with most factors having information gain when incorporating sequence features beyond short motifs (**Fig. S7C**). We find that nearly all RBPs have strong secondary structure preferences and ~50% of RBPs favor noncontiguous patterns. While the most common representation of RBP binding sites is a single position weight matrix (PWM), our data suggest that in many cases RBP specificity may be better described by pairs of short PWMs with variable spacing, and sometimes by PWMs describing structural preferences. We hypothesize that these subtle differences may be general features that allow RBPs that bind very similar motifs to select distinct targets in cells, a paradigm that has been put forth for pairs of RBPs (Smith et al., 2013).

In assaying a number of RBPs containing RRM, KH, and zinc finger domains, we noted several commonalities across RBPs with similar RBDs. For example, RRMs and ZFs bound to motifs rich in all four mononucleotides while KH domains did not bind any G-rich motifs, in agreement with previous crystallographic studies showing that G-recognition by KH domains is rare (reviewed by (Nicastro et al., 2015)). Further, while many motif clusters included RBPs from all three domain types (**Fig. 2A**), *k*mer enrichments were more highly correlated among RBPs with the same RBD type relative to RBPs with different RBD types, even after excluding paralogs (**Fig. S7F**). This observation suggests that RBPs with the same fold still harbor traces of a shared evolutionary history in RNA target recognition. We also noted domain-specific trends among preferences for contextual features. For example, while most RBPs overwhelmingly preferred unstructured motifs both in our assay (**Fig. 4A**) and *in vivo* (**Fig. S5B**), ZFs bound motifs encompassing a wide range of secondary structure contexts including those with greater stem content (**Fig. S7G**), and RBPs containing ZFs showed greater variability in their *P*_paired_ preferences than did RBPs without ZFs (**Fig. S7H**). These observations are consistent with a recent study finding that more than 20 ZF-containing proteins selectively bind highly-structured pre-miRNAs (Treiber et al., 2017). Proteins with KH domains in turn shared numerous properties, including a preference for binding to large hairpin loops, flanking nucleotide context preferences and the ability to bind to spaced motifs. Notably, all but one KH-containing RBP (SF1) have more than one KH domain or known homodimerization domains, suggesting that RBPs with KH domains may show a greater extent of RNA-binding cooperativity compared to other domain types.

RBP target selection in cells can be modulated by sequence and/or structure preference, but also by differential subcellular localization, expression, post-translational modifications, protein isoforms, and protein-protein interactions. Ultimately, all of these factors are likely to work together to specify post-transcriptional gene regulatory programs.

## SUPPLEMENTARY FIGURE LEGENDS

**Figure S1. RBNS assay and comparison to RNAcompete, Related to Figure 1**.

**A.** Histogram of the number of unique reads sequenced in all RBNS Input and Pulldown libraries, with the ~250,000 RNAcompete oligo sequences marked for comparison.

**B.** 10^th^ -90th percentile range of the mean base pairing probability (*P*_paired_) for all 5mer occurrences (averaged over all five positions) in RBNS input libraries and the RNAcompete custom array. The 1,024 5mers are sorted along the x-axis by increasing median RBNS *P*_paired_.

**C.** Dendrogram of clustered RBNS (red) and RNAcompete (black) experiments for the 31 RBPs that were assayed by both methods. Clustering was performed via the hclust function in R (with method=“average”) on the pairwise Pearson correlations of 5mer *R* Z-scores, for the set of 306 5mers that were enriched (Z≥3) in any of the 62 experiments.

**D.** Histogram of Pearson correlations of RBNS and RNAcompete for the 31 RBPs assayed by both methods as described in **C**.

**Figure S2. Overlapping specificities of RBPs, Related to Figure 2**.

**A.** Number of unique top 6mers among random subsamplings of the RBNS experiments versus randomly selected 6mers (similar to **Fig. 2C**), for the subset of 78 RBNS experiments that excludes any paralogs (left) or any RBPs that share at least 40% identity among any RBDs (right).

**B.** Similar to **Fig. 2C** and **Fig. S2A**, but for the top 15 6mers of each RBNS experiment instead of just the top 6mer. Black line determined from sets of 15 6mers in which the top 6mer was chosen at random, with the remaining 14 6mers matching the Hamming distance/shifts relative to the top 6mer observed among actual RBNS experiments (**Methods**).

**C.** Mapping of the four nucleotide frequencies in motif logos onto a 2D simplex, for both the actual 78 RBNS motifs (left), simulated RBNS motifs (center), and the resulting enrichment of RBNS versus simulated motifs in each simplex partition (right) (**Methods**). Gray boxes denote that none of the 78 RBNS motif frequencies mapped to that partition. Significance along margins of the enrichment simplex was determined by bootstrap Z-score (number of asterisks = Z-score). The data in the top and bottom rows is the same, with the A and C corners of the simplex switched so that each of the six dinucleotide combinations (AC, AG, AU, CG, CU, GU) is included along an edge in at least one of the two mappings. Upper right simplex is the same as **Fig. 2F**.

**Figure S3. RBNS motifs in the transcriptome and RBNS-derived splicing and stability RNA maps, Related to** **Figure 3**.

**A.** From left to right: Dendrogram of hierarchical clustering of RBPs by sequence logo similarity and 15 demarcated clusters by branch length cutoff (dashed line); protein name; top motif logo for each protein (up to here same as in **Fig. 2A**); schematic heat map representation of motif frequency (top half) and conservation (bottom half) within indicated human transcript regions; filled boxes indicating whether ≥3 of the top 15 RBNS 6mers for each protein overlap with splicing regulatory elements and 3’ UTR regulatory elements. Figure key at bottom.

**B.** Summary of RBNS regulatory activity inferred by enrichment of the top 10 RBNS 5mers in skipped exons (SEs) and flanking introns of exons significantly included or excluded in RNA-seq after RBP knockdown. Left: All RBPs assayed by both RBNS and KD/RNA-seq, ordered from top to bottom by RBNS motif (same ordering as in **Fig. 2A**), with KD in HepG2 and/or K562 denoted by orange/yellow bars. To the right: the top RBNS motif logo as in **Fig. 2A**, follow by a bar denoting whether there were significantly more SEs included upon KD (pink) or excluded upon KD (blue), where significance was determined by binomial proportion test *P*<0.05 with at least a 60%/40% bias (total number of SEs changing in each direction upon KD in bar plot on far right). Center: The inferred RBNS regulatory activity of the RBP over the SE and upstream and downstream 250nt of flanking intron, with strength of RBNS splicing regulatory activity of significant regions noted by color according to the green (ESE/ISE, significantly greater RBNS motif density for exons excluded upon KD) or red (ESS/ISS, significantly greater RBNS motif density for exons included upon KD) heat map. Exonic regions deemed significant if at least 20 of their 100 positions had significantly increased RNBS density (example for RBM25 in HepG2 and its significant positions for each direction of SEs shown in **Fig. 3C**), and upstream/downstream introns were deemed significant if 50 of their 250 intronic positions had significantly increased RNBS density. If an RBP was deemed to have significant RBNS regulatory activity in that region, the ratios of log_2_(RBNS density over changing SEs/RBNS density over control SEs) of all significant positions in that region were summed, and the maximum value was normalized to 1 over all RBPs.

**C.** Quantification of the proportion of intronic and exonic regions that show significant enrichment of RBNS motifs that enhance (top) or silence (bottom) splicing in **B**.

**D.** Similar to **B**, but for inferred RBNS regulatory activity on gene expression levels based on RBNS motif density in the 3’ UTRs of genes significantly up-or down-regulated upon KD. Each RBP was deemed to have significant RBNS regulatory activity if 10 of the 100 positions of the meta-3’ UTR had increased RBNS density as shown for the TAF15 example in **Fig. 3F** (Destab.=increased RBNS density in 3’ UTRs of genes up-regulated upon KD; Stab.=increased RBNS density in 3’ UTRs of genes down-regulated upon KD).

**E.** Similar to **Fig. 3F**, but for SRSF5 KD in K562 (left) and HepG2 (right) cells.

**Figure S4. Conservation of RBNS motifs in the transcriptome and *in vivo* binding of RBPs, Related to Figure 3.**

**A.** Cumulative distribution of phyloP conservation Z-scores for RBNS 5mers and non-RBNS 5mers in exons and upstream and downstream flanking introns. Distributions are shown separately for constitutive, alternative, and ancient alternatively spliced exons.

**B.** Comparison of RBNS and eCLIP motifs for RBPs assayed by both techniques (reproduced with permission from (Van Nostrand et al., 2017)). The top RBNS and eCLIP motif are shown for each RBP (**Methods**), clustered by RBNS motif as in **Fig. 2A**. 17 of the 26 RBPs with significant overlap in the 5mers comprising the RBNS and eCLIP logos (*P*<0.05, hypergeometric test) marked with a star to the right of the eCLIP logo.

**C.** Left: Enrichment of RBNS 5mers (averaged among all RBNS 5mers with Z-score≥3) in eCLIP peaks that occur in both replicates (peaks overlap by at least 1 position) relative to peaks that occur in only one replicate. Right: Enrichment of RBNS 5mers (averaged among all RBNS 5mers with Z-score≥3) in eCLIP peaks that occur in both HepG2 and K562 (peaks overlap by at least 1 position) relative to peaks that occur in only one cell type.

**D.** Genome browser snapshot of a portion of the 3’ UTR of the *STOM* gene and eCLIP densities (normalized to input) for TIA1 and HNRNPC bound to two different locations in this region. Tracks show instances of U_5_ motifs in this region (red, top) and eCLIP peaks are marked in black below each density track. Motifs derived from RBNS and eCLIP peaks for each RBP shown on the left.

**Figure S5. Distribution of enrichments along reads and *in vivo* structural preferences of RBPs, Related to Figure 4**.

**A.** The frequency of the top RBNS 6mer at each position of the random region, relative to a uniform distribution at all positions (RBPs with random 20mers, top; random 40mers, bottom). Difference from uniform denoted by the green heat map bar to the right, calculated as the KL-divergence of the observed frequency at each position relative to a uniform distribution; RBPs are sorted by decreasing divergence. RBPs and their top 6mers noted on left, with ZNF326 and RBM22 marked as the 1^st^ and 3^rd^ most unequal distributions across 20mers, respectively.

**B.** Correlation of *R* value profiles of the top 5mer across five bins of increasing P_paired_ for RBPs assayed by both RBNS (x-axis) and eCLIP (y-axis). For each assay, *R* was calculated in each of the 5 *P*_paired_ bins and the monotonicity of *R* over the 5 bins was calculated (-10 monotonicity = 5 bins monotonically decreasing *R* with increasing *P*_paired;_ 10 monotonicity = 5 bins monotonically increasing *R* with increasing *P*_paired_).

**Figure S6. Bipartite core spacing, flanking nucleotide context, and degenerate pattern binding preferences, Related to Figure 5**.

**A.** Enrichment as a function of spacing in 0nM protein controls. Top 10 pairs of 3mers are shown for each experiment at each spacing.

**B.** Relative preference for spacing (ΔF, **Methods**) grouped by whether RBPs have a single RBD, multiple RBDs, or show a potential to multimerize (based on literature references and RBP regions cloned in our assay).

**C.** Enrichment as a function for spacing for the RBPs that show an increase in enrichment as a function of spacing in **Fig. 5C**. Experiments done with protein concentrations that are lower or equal to the concentration with maximum enrichment are shown.

**D.** Flanking nucleotide context preferences for RBPs with KH domains, with and without controls for RNA secondary structure. “Match pPaired” represents motif occurrences that are sampled to have the same mean base pairing probability in the input and pulldown libraries. “Motifs in hairpins” represents motif occurrences that have the 5-letter code “HHHHH” in the minimum free energy structure in both the input and pulldown libraries.

**E.** Enrichment for particular nucleotides in reads with no high-affinity motifs for FUS (left), IGF2BP1 (right).

**F.** Enrichments for degenerate patterns of length 12 with 6 fixed bases shown for HNRNPK (left), RBFOX2 (right). Only the top 5% of degenerate patterns are shown and the red lines indicate the enrichment of the linear 6mers. For HNRNPK, degenerate patterns where 5 out of the 6 fixed positions are Cs are in blue.

**G.** Enrichment of HNRNPK motifs in eCLIP data. Linear: top 10 6mers; bipartite: top 10 spaced 6mers; context: top 10 degenerate *k*mers with 6 Cs (those shown in **Fig. 5H** left).

**H.** Bar plot of enrichments for top 10 degenerate *k*mers of length 12 with 6 Ns for CELF1 (top) and FUBP1 (bottom).

**Figure S7. Sequence context effects on RBP binding, Related to Figure 6**.

**A.** Example of the distance between feature-specific *R* value profiles for PCBP2 and RBM23 for *^P^*_paired._

**B.** Dispersal of specificities as in **Fig. 6B** for all other clusters.

**C.** Top: The maximum *R* value (the greater of the best linear or bipartite motif) for each RBP. Bottom: The relative entropy of the distribution of pulldown vs. input top 6mer occurrences in five context bins for each of four context features (*P*_paired_, *P*_flank_, core spacing and flanking nucleotide context). The top 6mer used for each of the 15 clusters marked to the left of each cluster.

**D.** Distribution of UAAUUA motif counts among pulldown reads for cluster 1 proteins for three context features (*P*_paired_, *P*_flank_ and flanking nucleotide context) used to calculate the relative entropy in **Fig. 6C**. *P*_paired_ and *P*_flank_ bins were set empirically for each RBP such that 20% of input UAAUUA occurrences were in each bin; nucleotide context bins were set as described in Methods. The 7 RBPs were clustered (dendrogram shown on left) according to their context preferences over the entire 15 different context bins.

**E.** Tissue specificity of gene expression profile in 40 human tissues from the GTEx consortium for RBPs within a motif cluster versus those unclustered. *P*-value determined by Wilcoxon rank-sum test.

**F.** Distribution of Pearson correlations of the *R* Z-scores of top 5mers for RBPs within the same motif cluster, separated by RBP pairs that have different RBD types (left) or the same RBD type (all RBPs: right; no paralogs: center). The *R* Z-score of the top 18 5mers for each RBP pair were used (the median number of enriched 5mers), and RBPs with only 1 RBD type were included. \*\**P*<0.05 by Wilcoxon rank-sum test.

**G.** Increase in stem content (*f*_stem,PD_ × log_2_(*f*_stem,PD_/*f*_stem,IN_), where *f*_stem,PD_ is shown for RBM22 and ZNF326 in **Fig. 4E,F**) averaged over the top 6mer of each RBP, separated by RBPs that do vs. do not contain a ZF. *P*-value determined by Wilcoxon rank-sum test.

**H.** Comparison of differences in *P*_paired_ preferences among pairs of RBPs (|*P*_paired_ ratio,RBP_1_ – *P*_paired_ ratio,RBP_2_|, where *P*_paired_ ratio is *P*_paired, PD_/*P*_paired, IN_ averaged over the top 6mer as shown in the right bar of **Fig. 4A**), separated by RBPs that do vs. do not contain a ZF. *P*-value determined by Wilcoxon rank-sum test.

## SUPPLEMENTARY TABLES

**Table S1.** Number of reads in all RBNS experiments

**Table S2.** List of primers and cloned regions for all assayed RBPs

**Table S3.** Summary of RBNS experiments including experimental procedures and derived motifs.

**Table S4:** *P*_paired_ of each position in input and pulldown library, recalculated *R* by *P*_paired_ bin, and recalculated *R* values by structural element for top 6mer of each RBNS

**Table S5:** Bipartite motifs and nucleotide context preferences for all assayed RBPs.

**Table S6:** Motif clusters and their significantly divergent features.

## ACKNOWLEDGEMENTS

This work was funded by the National Human Genome Research Institute ENCODE Project, contract U54HG007005, to B.R.G. (principal investigator) and C.B.B., and G. W.Y. (co-principal investigators). We thank Dr. Timothy Hughes for kindly providing bacterial expression constructs for RBPs.

The 78 RBNS data sets described here can be obtained from the ENCODE project website at https://www.encodeproject.org/search/?type=Experiment&assay_title=RNA+Bind-N-Seq&assay_title=RNA+Bind-n-Seq and via the Accession Numbers in Table S3. The authors declare no competing financial interests. Correspondence and requests for materials should be addressed to Christopher B. Burge.

## METHODS

### Cloning of RNA binding protein domains

In most cases, RBPs were selected from a curated set of high-confidence annotations consisting of factors with well-defined RNA binding domains or with previous experimental evidence of RNA binding (Van Nostrand et al., 2017). Regions of each protein containing all RBDs plus ~50 amino acids flanking the RBD were cloned into the pGEX6 bacterial expression construct (GE Healthcare). A list of all constructs generated and primer sequences used is shown in **Table S2**.

### Bacterial expression and protein purification

Transformed Rosetta Cells (Novagen) were cultured in SuperBroth until optical density reached 0.6, cultures were transferred to 4° C and allowed to cool. Protein expression was induced for 14-20 hrs with IPTG at 15° C. Cells were pelleted, lysed (Qproteome Bacterial Protein Prep Kit, Qiagen) for 30 min in the presence of protease inhibitor cocktail (Roche), sonicated and clarified by centrifuging at >8,000 rpm, passed through a .45 μM filter (GE) and purified using GST-sepharose in either column format (GST-trap FF, GE) or 96-well format (GSTrap 96-well Protein Purification Kit, GE). Generally, 250mL bacterial cultures used for column purifications and 50mL for 96-well plate purifications (note: 8 wells of a 96-well plate were used per protein so that up to 12 proteins were purified per plate at a time). Eluted proteins were concentrated by centrifugation (Amicon Ultra-4 Centrifugal Filter Units) and subjected to buffer exchange (Zeba Spin Desalting Columns, 7K MWCO, Life Technologies) into Final Buffer (20mM Tris pH 7, 300mM KCl, 1M DTT, 5mM EDTA, 10% glycerol). Proteins were quantified using Bradford Reagent (Life Technologies) and purity and quality of protein was assessed by PAGE followed by Coomassie staining (all gels are available at https://www.encodeproject.org/search/?type=Experiment&assaytitle=RNA+Bind-N-Seq&assaytitle=RNA+Bind-n-Seq).

### Production of random RNAs by *in vitro* transcription

Single-stranded DNA oligonucleotide and random template were synthesized (Integrated DNA Technologies) and gel-purified as previously described (Lambert et al., 2014). Synthesis of random region of the template DNA oligo was “hand-mixed” to achieve balanced base composition. An oligo matching T7 promoter sequence was annealed to the random template oligo by mixing in equal parts bringing to 70° C for 2 min and allowing to cool by placing at room temperature. T7 Template: 5’ CCTTGACACCCGAGAATTCCA(N)_20_GATCGTCGGACTGTAGAACTCCCTATAGTGAG TCGTA T7 oligo: 5’ TAAT ACGACT CACTAT AG GG RNA was synthesized by transcribing 6uL of 25uM annealed template and T7 oligo in a 100 μL reaction (Hi-Scribe T7 transcription kit, NEB) according to manufacturer’s protocol) or with a custom protocol usting T7 polymerase (NEB) for larger-scale preps. RNAs were then DNAse-treated with RQ1 (Promega) and subjected to phenol-chloroform extraction. RNA was suspended in nuclease free water and resolved on a 6% TBE-Urea gel (Life Technologies). RNA was excised and gel-extracted as previously reported (Lambert et al., 2014). RNA was aliquoted and stored at −80° C.

### RNA Bind-n-Seq Assay

All steps of the following binding assay were carried out at 4° C. Dynabeads MyOne Streptavidin T1 (Thermo) were washed 3X in binding buffer (25mM tris pH 7.5, 150 mM KCl, 3mM MgCl2, 0.01% tween, 500 ug/mL BSA, 1 mM DTT). 60 uL of beads per individual protein reaction were used. 60 uL RBP diluted (see below for protein concentrations used) in binding buffer were allowed to equilibrate for 30 minutes at 4°C in the presence of 60 uL of washed Dynabeads MyOne Streptavidin T1. After 30 min of incubation, 60 uL of random RNA diluted in binding buffer was added bringing the total reaction volume to 180 uL. The final concentration per reaction of each of the components was 1uM RNA; 5, 20, 80, 320 or 1300 nM of RBP; and 60uL of Dynabeads MyOne Streptavidin T1 stock slurry washed and prepared in binding buffer. Each reaction was carried out in a single well of a 96-well plate. After 1 hr, RBP-RNA complexes were isolated by placing 96-well plate on a magnetic stand for 2 min. Unbound RNA was carefully removed from each well and the bound RNA complexes were washed with 100 uL of wash buffer (25mM tris pH 7.5, 150 mM KCl, 0.5 mM EDTA, 0.01% tween). Immediately after adding wash buffer the plate was placed on the magnet and wash was removed after ~1min. This procedure was repeated 3 times. RBP-RNA complexes were eluted from Dynabeads MyOne Streptavidin T1 by incubating reaction at room temperature for 15 minutes in 25 uL of elution buffer (4mM biotin, 1× PBS), the eluate was collected, the elution step was repeated, and eluates were pooled. RNA was purified from elution mixture by adding 40 uL AMPure Beads RNAClean XP (Agencourt) beads and 90 uL of isopropanol and incubating for 5 minutes. 96-well plate was placed on a magnetic stand and supernatant was discarded. Beads were washed 2X with 80% ethanol, dried, and RNA was eluted in 15 uL of nuclease-free water. The extracted RNA was reverse transcribed into cDNA with Superscript III (Invitrogen) according to manufacturer’s instructions using the RBNS RT primer. To prepare the input random library for sequencing, 0.5 pmol of the RBNS input RNA pool was also reverse transcribed. To make Illumina sequencing libraries, primers with Illumina adapters and sequencing barcodes were used to amplify the cDNA by PCR using Phusion DNA Polymerase (NEB) with 10-14 PCR cycles. PCR primers always included RNA PCR 1 (RP1) and one of the indexed primers as previously reported (Lambert et al., 2014). PCR products were then gel-purified from 3% agarose gels and quantified and assessed for quality on the Bioanalyzer (Agilent). Sequencing libraries for all concentrations of a the RBP as well as the input library were pooled in a single lane and sequenced on an Illumina HiSeq2000 instrument.

### IRNA Bind-n-Seq data processing and motif logo generation

RBNS Amer enrichments (*R* values) were calculated as the frequency of each Amer in the pulldown library reads divided by its frequency in the input library; enrichments from the pulldown library with the greatest enrichment were used for all analyses of each respective RBP. Mean and standard deviation of *R* values were calculated across all 4^*k*^ *k*mers for a given *k* to calculate the RBNS Z-score for each *k*mer.

RBNS motif logos were made from following iterative procedure on the most enriched pulldown library for *k*=5: the most enriched kmer was given a weight equal to its enrichment over the input library (=*R*-1), and all occurrences of that *k*mer were masked in both the pulldown and input libraries so that stepwise enrichments of subsequent Amers could be used to eliminate subsequent double counting of lower-affinity ‘shadow’ *k*mers (e.g., only GGGGA occurrences not overlapping a higher-affinity GGGGG would count towards its stepwise enrichment). All enrichments were then recalculated on the masked read sets to obtain the resulting most enriched *k*mer and its corresponding weight (=stepwise *R*-1), with this process continuing until the enrichment Z-score (calculated from the original *R* values) was less than 3. All *k*mers determined from this procedure were aligned to minimize mismatches to the most enriched *k*mer, with a new motif started if the *k*mer could not be aligned to the most enriched *k*mer in one of the following 4 ways: one offset w/ 0 mismatches (among the 4 overlapping positions); 1 offset w/ 1 mismatch; no offset w/ 1 mismatch; 2 offsets w/ 0 mismatches. The frequencies of each nucleotide in the position weight matrix, as well as the overall percentage of each motif, were determined from the weights of the individual aligned kmers that went into that motif; empty unaligned positions before or after each aligned kmer were given pseudocounts of 25% each nucleotide, and outermost positions of the motif logo were trimmed if they had had unaligned total weight >75%. To improve the robustness of the motif logos, the pulldown and input reads were each divided in half and the above procedure was performed independently on each half; only *k*mers identified in corresponding motif logos from both halves were included in the alignments to make the final motif logo (weight of each Amer averaged between the two halves). In **Fig. 2A**, only the top RBNS motif logo is shown if there were multiple (all motifs displayed on the ENCODE portal within the “Documents” box of each experiment, with the proportion of each motif logo determined by computing the relative proportion of each motif’s composite Amer weights). Motif logos were made from the resulting PWMs with Weblogo 2.0 (Crooks et al., 2004). In addition to those displayed for 5mers with a Z-score=3 cutoff, for comparison motif logos were also made using: 5mers with Z-score=2 cutoff, 6mers with Z-score=2 cutoff, and 6mers with Z-score=3 cutoff; additionally, different rules of when to start a new logo vs. add to an existing one were tried. Logos for 5mers with Z-score=3 cutoff and the rule mentioned above appeared to strike the best balance of capturing a sufficient number of Amers to accurately represent the full spectrum of the RBP’s binding specificity but did not create a number of secondary, largely similar motifs and thus were chosen to use across all 78 RBPs.

The RBNS pipeline is available at: https://bitbucket.org/pfreese/rbnspipeline

### Clustering of RBNS motifs

A Jensen-Shannon divergence (JSD)-based similarity score between each pair of top RBNS motif logos was computed by summing the score of the *j* overlapping positions between RBP A and RBP B:

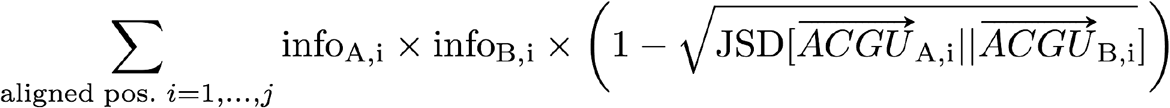

where info_A,i_ is the information content in bits of motif A at position *i* and ACGU_A,i_ is the vector of motif A frequencies at position *i* (vectors sum to 1.)

This score rewards positions with higher information content (scaled from positions positions with 100% one nucleotide given maximum weight to degenerate positions with 25% each nucleotide given zero weight) and more aligned positions (more positions *j* contributing to the summed score).

This similarity score was computed for each possible overlap of the two logos (subject to at least four positions overlapping, i.e., *j*≥4), and the top score with its corresponding alignment offset was used. The matrix of these scores were normalized to the maximum score over all RBP pairs and clustered using the linkage function with centroid method in scipy.cluster.hierarchy to obtain the dendrogram shown in **Fig. 2A**, with the 15 RBP groupings derived from a manually-set branch length cutoff. This branch length cutoff was chosen to balance the competing interests of maximizing the number of paralogous proteins within the same cluster (more stringent cutoffs eliminated PCBP4 from the cluster containing PCBP1 and PCBP2; it also did not include RBM4 and RBM4B to be in a cluster) and minimizing differences between primary motifs within the same cluster (less stringent cutoffs included the distinct UAG-containing MSI1/UNK/HNRNPA0 motifs within the same cluster as the AU-rich RBPs, for example).

### Comparison with RNAcompete

5mer scores were derived from publicly available 7mer Z-scores by computing the mean across all 7mers containing a given 5mer (http://hugheslab.ccbr.utoronto.ca/supplementary-data/RNAcompeteeukarya/). Correlations between RBNS and RNA-compete experiments were computed by taking the Pearson correlation of Z-scores for all 5mers which had a Z-score ≥ 3 for at least one of the 31 RBPs in common between both assays.

### Analysis of RBNS motif frequency and conservation in the transcriptome

Frequency and conservation of an RBP’s motif (**Fig. 2A**) were calculated for each RBP by taking a weighted average of all its motif logo 5mers as reported in **Table S3**.

For each protein-coding gene, the most highly expressed Gencode version 19 transcript in HepG2 or K562 cells was used to determine 5’ and 3’UTR regions. Exon sequences were all internal exons between the 5’ and 3’UTRs. The upstream intron sequences were −130 to −30 of each of these exons and the downstream intron sequences were +9 to 109 (i.e., 100 nt regions flanking internal exons excluding the splice sites). The frequency of all 1,024 5mers was computed in each of these regions, and the frequency shown in **Fig. S3A** is: (RBP’s weighted average frequency in the region) × 4^5^.

Conservation of all 1,024 5mers was computed in each of the aforementioned regions by taking the average 46-way phyloP conservation over the 5mer. The mean and standard deviation (over the 1,024 5mers) was computed separately in each region, and the conservation shown in **Fig. S3A** is the Z-score of the RBP’s weighted conservation in the region.

### Overlap of RBNS 6mers with splicing and stability regulatory elements

Splicing regulatory elements were taken from: ESS and ESE: (Ke et al., 2011) and (Rosenberg et al., 2015); ISE: (Wang et al., 2012); ISS: (Wang et al., 2013). For classification of proteins as overlapping we first generated a set of ISS, ISE, ESS, or ESE 6mers. 6mers derived from 10mers screened by ISE: (Wang et al., 2012); ISS: (Wang et al., 2013) were used as ISE and ISS elements. For ESE and ESS elements, the top 400 ESS and ESE reported in the Supplementary Table 1 of (Ke et al., 2011) (“ESRseqScore”) and the top 400 ESS and ESE derived from the 5’splice site selection reported by (Rosenberg et al., 2015) as indicated in their manuscript were combined. A protein was classified as binding to ISSs, ISEs, ESSs, or ESEs if 3 of its top 15 6mers overlapped the ISS, ISE, ESS or ESE 6mers (As is shown in **Fig. S3A**, black and white boxes). For analysis of 6mer splicing regulatory scores (**Fig. 3B**) the reported “ESRseqScores” from (Ke et al., 2011) were used.

3’UTR regulatory 6mers were derived from (Oikonomou et al., 2014). Only 6mers with ≥100 occurrences across all designed sequences were used (totaling 1303 6mers) in order to derive a mean 6mer score with sufficient coverage in different contexts. 6mer repressor and activator scores were obtained by averaging scores (log2 frequency as described in the original manuscript) across all oligos containing that 6mer in the low (L10) and high (H10) Dual-reporter Intensity Ratio bins, respectively. Activator and repressor scores were averaged across both replicates (Libraries A and B). 6mers with an overall score ≥ 0.25 were used, where regulatory score=|log_2_(repressor score)-log_2_(activator score)|.

### Analysis of eCLIP for motif discovery, regulation and overlapping targets

eCLIP datasets were produced by the Yeo Lab through the ENCODE RBP Project and are available at https://www.encodeproject.org/search/?type=Experiment&assay_title=eCLIP.

For all analyses, only eCLIP peaks with an enrichment over input ≥ 2 were used. Peaks were also extended 50 nucleotides in the 5’ direction as the 5’ start of the peak is predicted to correspond to the site of crosslink between the RBP and the RNA.

To produce eCLIP logos in a similar manner for comparison with RBNS logos, an analogous procedure to creating the RBNS motif logos was carried out on the eCLIP peak sequences: the two halves of the RBNS pulldown reads were replaced with the two eCLIP replicate peak sequences, and the input RBNS sequences were replaced by random regions within the same gene for each peak that preserved peak length and transcript region (5’ and 3’ UTR peaks were chosen randomly within that region; intronic and CDS peaks were shuffled to a position within the same gene that preserved the peak start’s distance to the closest intron/exon boundary to match sequence biases resulting from CDS and splice site constraints). The enrichment Z-score threshold for 5mers included in eCLIP logos was 2.8, as this threshold produced eCLIP logos containing the most similar number of 5mers to that of the Z=3 5mer RBNS logos. Each eCLIP motif logo was filtered to include only 5mers that occurred in both corresponding eCLIP replicate logos. eCLIP motif logos were made separately for all eCLIP peaks, only 3’UTR peaks, only CDS peaks, and only intronic peaks, with the eCLIP logo of those 4 (or 8 if CLIP was performed in both cell types) with highest similarity score to the RBNS logo shown in **Fig. S4B**, where the similarity score was the same as previously described to cluster RBNS logos. To determine overlap significance of RBNS and eCLIP, a hypergeometric test was performed with the 5mers in all (not just the top) logos for: RBNS logo 5mers, eCLIP logo 5mers (for peaks in the region with highest similiarity score to the RBNS logo), and 5mers in their intersection among the background of all 1,024 5mers; overlap was deemed significant if P<0.05.

All eCLIP/RBNS comparisons were for the same RBP with the following exceptions in which the eCLIP RBP was compared to its paralogous RBNS protein: KHDRBS2 (KHDRBS1 RBNS); PABPN1 (PABPN1L RBNS); PTBP1 (PTBP3 RBNS); PUM2 (PUM1 RBNS); and RBM15 (RBM15B RBNS).

For **Fig. 3G**, the Pearson correlation between eCLIP experiments was assessed by computing the mean eCLIP coverage across 3’UTRs of all genes. 3’UTRs were split into windows of ~100 nucleotides and the mean base-wise coverage (eCLIP coverage divided by input coverage) was calculated in each window. Pairs of RBPs were assigned as paralogs according to their classification in Ensembl. Pairs of RBPs were assigned as having overlapping motifs if at least 2 of their 5 top 5mers overlapped; RBPs with specificities determined from RBNS and RNAcompete were pooled (Ray et al., 2013).

### Analysis of RNA-seq datasets for regulation and RBNS Expression & Splicing Maps

RNA-seq after shRNA knockdowns of individual RBPs in HepG2 and K562 cells (two KD and two control RNA-seq samples per RBP) were produced by the Graveley Lab through the ENCODE RBP Project and are available at: https://www.encodeproject.org/search/?type=Experiment&assaytitle=shRNA+RNA-seq.

Splicing changes upon KD were quantified with MATS (Shen et al., 2012), considering only skipped exons (SEs) with at least 10 inclusion + exclusion junction-spanning reads and a ψ between 0.05 and 0.95 in the averaged control and/or KD samples. SEs that shared a 5’ or 3’ splice site with another SE (i.e., those that are part of an annotated A3’SS, A5’SS, or Retained Intron) were eliminated. If multiple pairs of upstream & downstream flanking exons were quantified for an SE, only the event with the greatest number of junction-spanning reads was used. SEs significantly excluded or included upon KD were defined as those with a P-value < 0.05 and |Δψ| ≥ 0.05. Control SEs upon KD were those with a P-value=1 and |Δψ| ≤ 0.02.

Differentially expressed genes upon KD were called from DEseq2 (Love et al., 2014), considering genes that had a ‘baseMean’ coverage of at least 1.0 and an adjusted P-value < 0.05 and |log_2_(FC)| ≥ 0.58 (1.5-fold up or down upon KD). Candidate control genes upon KD were taken from those with a P-value > 0.5 and |log_2_(FC)| ≤ 0.15; from this set of genes, a subset matched to the deciles of control (i.e., before KD) gene expression levels of the differentially expressed genes was used. The last 50nt of each gene’s ORF and 3’UTR sequence were taken from the Gencode version 19 transcript with the highest expression in the relevant cell type (HepG2 or K562).

‘RBNS splicing maps’ were made by taking the three sets of SEs included, excluded, or control upon KD and extracting their exonic and upstream/downstream flanking 250nt sequences. At each position of each event, it was determined whether the position overlapped with one of the top 10 RBNS 5mers for that RBP in any of the five registers overlapping the position. Then to determine if the RBNS density was significantly higher or lower for included/excluded SEs at a position relative to control SEs at that position, the number of positions in a 20bp window on each side (total 41 positions) covered by RBNS motifs was determined for each of the events, with significance determined by P-value<0.05 in a Wilcoxon rank-sum test on the control vs. changed events in the desired direction upon KD. Exonic regions were deemed to have ESE or ESS RBNS regulatory activity if 20 of the 100 exonic positions among SEs excluded or included upon KD, respectively, had significantly higher RBNS motif coverage than control SEs. The upstream and downstream intronic regions were each individually deemed as ISE or ISS regions if 50 of the 250 intronic positions had significantly higher RBNS motif coverage.

‘RBNS stability maps’ were made in an analogous manner, but for genes up-or down-regulated compared to control genes upon KD. The 3’UTR sequence was divided into 100 segments of roughly equal length and the proportion of positions covered by RBNS motifs in each segment were used for each bin of the meta-3’UTR. An RBP was deemed to have significant RBNS regulatory activity if 10 of the 100 positions of the meta-3’UTR for up-or down-regulated genes had increased RBNS density relative to control genes.

### Generation of random sets of ranked 6mer lists with edit distances to top 6mer matching RBNS

Because the ranked lists of top enriched *k*mers (e.g., the top 15 6mers) are highly constrained depending on what the most enriched *k*mer is (e.g., 6mers 2-15 are typically Hamming distance of 1 and/or shifted by 1 from the top 6mer), as background sets for comparison to actual RBNS 6mer lists we sought to create groups of 6mers that matched the observed RBNS patterns of Hamming distances and shifts from the top 6mer for any given randomly selected *k*mer. To do this, for each of the 78 RBNS experiments we first calculated the edit distance from 6mer_1_ to 6mer_1_, where 6meri is the most enriched 6mer and *i*=2, …, 15 is the *i*th enriched 6mer (e.g., 6mer_8_ might have a mismatch at position two compared to 6mer_1_ and then be shifted to the right by 1 position). Then, for all 4,096 starting 6mers, we created 78 ranked lists of 15 6mers, each of which matched the observed edit distances to the top 15 list of an actual RBNS experiment. The expected number of network edges in **Fig. 2B**, and the ‘random’ number of edges in **Fig. 2D** were performed by selecting random lists from these 4,096*78 possibilities.

### RBNS RBP groups without paralogs or RBPs with any RBD pair sharing 40% identity

#### No Paralogs (*n*=52)

A1CF, BOLL, CELF1, CNOT4, CPEB1, DAZ3, EIF4G2, ELAVL4, ESRP1, EWSR1, FUBP1, HNRNPA2B1, HNRNPC, HNRNPK, HNRNPL, IGF2BP1, ILF2, MBNL1, NUPL2, PABPN1L, PRR3, PTBP3, PUM1, RBFOX2, RBM15B, RBM22, RBM23, RBM24, RBM25, RBM4, RBM41, RBM45, RBM47, RBM6, RBMS2, RC3H1, SF1, SFPQ, SNRPA, SRSF10, SRSF11, SRSF2, SRSF4, SRSF8, TARDBP, TIA1, TRA2A, TRNAU1AP, UNK, ZCRB1, ZFP36, ZNF326

#### No RBPs sharing >40% identity among any RBDs (*n*=47)

A1CF, BOLL, CELF1, CNOT4, CPEB1, EIF4G2, ELAVL4, EWSR1, FUBP3, HNRNPA0, HNRNPCL1, HNRNPDL, HNRNPH2, HNRNPL, IGF2BP1, ILF2, KHDRBS3, MBNL1, NOVA1, NUPL2, PABPN1L, PCBP2, PRR3, PTBP3, PUF60, PUM1, RBFOX3, RBM15B, RBM22, RBM24, RBM25, RBM41, RBM45, RBM4B, RBM6, RBMS2, SFPQ, SNRPA, SRSF11, SRSF8, SRSF9, TARDBP, TIA1, TRA2A, TRNAU1AP, ZFP36, ZNF326

Pairwise RBD alignments were performed using ClustalW2 (Larkin et al., 2007) and percent identities (as shown in **Fig. 1C** and **5D**) were calculated as the percentage of identical positions relative to the number of ungapped positions in the alignments.

### Network map of overlapping affinities

The lists of top 15 6mers for each RBP were intersected to get the number in common – those with 2 or more were deemed significant and connected by an edge (*P*<0.05 by Hypergeometric test, as well as by simulations based on the empirical distribution from random sets of ranked 6mer lists with edit distances to top 6mer matching RBNS as described above). The resulting network was visualized with Cytoscape (Shannon et al., 2003).

### Motif entropy analysis

To construct a set of ‘shuffled’ motifs that matches the overall nucleotide composition of the 78 RBNS motifs but removes any positional correlations within a motif, individual columns of each RBNS motif (including all motifs for an RBP if there was more than one) were pooled to be sampled from. A ‘shuffled’ motif was constructed by randomly sampling 5 or 6 columns (with probability 2/3 and 1/3, respectively, to roughly match the lengths of RBNS motifs) from this pool and concatenating them, repeated to construct 100,000 shuffled motifs.

The frequency of the four bases in each logo was calculated by averaging over all positions in the motif. This frequency vector (=[f_A_, f_C_, f_G_, f_U_], f_A_+f_C_+f_G_+f_U_=1) was mapped onto a square containing corners at [+/−1, +/−1] using two different orderings of the 4 corners, which together contain all 6 dinucleotide combinations (AC, AG, AU, CG, CU, GU) as edges:

1. Purine/Pyridine diagonals:

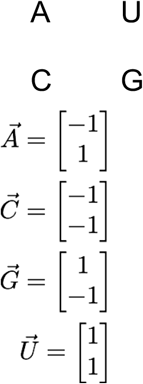
2. Purine/Pyridine edges:

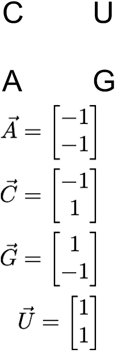

To map the frequency vector to its coordinates (u, v) within the unit circle, the frequency vector was normalized to the largest component:

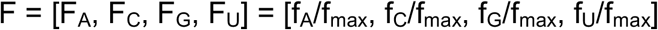
where f_max_ = max(f_A_, f_C_, f_G_, f_U_) 
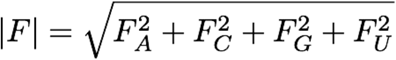

and (u, v) was computed as:

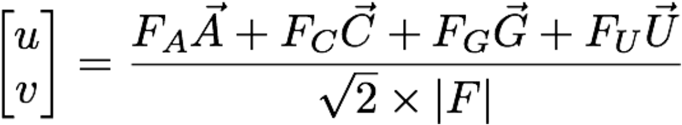

The elliptical grid mapping was used to convert the (u, v) coordinates within the unit circle to the corresponding position (x, y) within a square containing corners at [+/−1, +/− 1]:

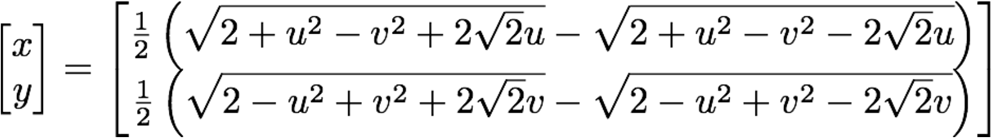

The simplex grid shown was divided into 11 equal parts along both dimensions, and the density in each of the 121 squares was computed for the 78 RBNS motifs and 100,000 shuffled motifs to get enrichments

To determine significance via bootstrapping, 1,000 different shuffled motif distributions over the grid were computed. In each of the 1,000 bootstraps, the 100,000 shuffled motifs were drawn from a different starting pool of motif columns: rather than all 78 RBPs’ motifs contributing once to the pool, a random sampling (with replacement) of the 78 RBPs was performed, and those motifs’ columns served as the starting pool for the 100,000 shuffled motifs. The mean and standard deviation of these 1,000 bootstraps were computed for each box, row, and column, and rows/columns of the grid for which the density of the 78 RBNS motif logos had a Z-score greater than 2 were marked (number of asterisks = Z-score, rounded down).

### RNA secondary structure analysis

The RNA base pairing probability was extracted from the partition function of RNAfold: “RNAfold ∐p --temp=X”, where X was 4° or 21°C depending on what temperature the binding reaction was conducted at (See Table S3) (Lorenz et al., 2011). For each pulldown library, reads were randomly selected to match the distribution of C+G content among input reads; all enrichments were recalculated for these C+G-matched pulldown reads for **Fig. 4** and **Fig. 6**. Reads were folded with the 5’ and 3’ adapters (24 and 21 nt, respectively), resulting in folded sequences of length 65 and 85 for 20mer and 40mer RBNS experiments, respectively.

Secondary structural element analyses were performed by using the forgi software package (Kerpedjiev et al., 2015). For each read, to mirror the partition function rather than relying solely on the Minimum Free Energy structure, 20 random suboptimal structures with probabilities equal to their Boltzmann weights were sampled and averaged over (“RNAsubopt --temp=X-stochBT=20”). In **Fig. 4D**, 6mers counting toward: ‘loop’ were: H_6_, M_6_, I_6_; ‘stem’ was S_6_; ‘bulged stem’ were 6mers matching the pattern SXXXXS, where XXXX contained 1-3 S.

Bin limits for the motif structure analyses (*P*_paired_) were: 0-0.2 (bin 1); 0.2-0.4 (bin 2); 0.4-0.6 (bin 3); 0.6-0.8 (bin 4); and 0.8-1.0 (bin 5). Bin limits for flanking structure analyses (Pflank) were: 0-0.3 (bin 1); 0.3-0.45 (bin 2); 0.45-0.6 (bin 3); 0.6-0.75 (bin 4); 0.75-1.0 (bin 5). *P*_paired_ was calculated as the average over the six positions of the 6mer; *P*_flank_ was calculated as the average over all other positions in the read. The continuous measures of preference for motif and flanking preference displayed in Fig. 6B,C were computed as:

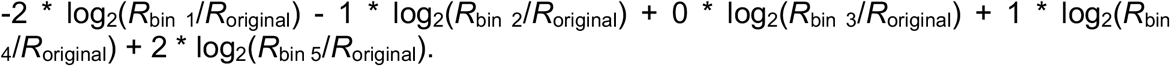

RBNS structure profiles were compared to eCLIP structure profiles in the region with the greatest number of eCLIP peaks. Bound RBNS motifs were selected from the transcriptome region that showed the highest enrichment for the number of peaks (5’UTR/3’UTR/introns/CDS). Motifs that were not bound were selected from the same gene regions as bound motifs and matched for the same genes. Motifs were folded with 50 nucleotides of flanking sequence on both sides using RNAfold (Lorenz et al., 2011). Motifs (both bound and not bound) were then binned by their mean base pairing probability (same bins as RBNS) and the fraction of bound motifs in each bin was then computed. The monotonicity of *R* over *P*_paired_ bins for RBNS and eCLIP was computed by taking all 10 comparisons over the 5 bins, adding 1 if *R* was greater in the higher *P*_paired_ bin or subtracting 1 if it was lower in the higher *P*_paired_ bin.

### Determination of bipartite motifs

Enrichments were computed for all pairs of the top 10 enriched 3mers, with a spacer of length i=0-10 (in total: 10*10*(i+1) combinations), where the enrichment was defined as the fraction of pulldown reads with a motif relative to the fraction of input reads with a motif. The enrichment for each spacing was computed as the mean enrichment of the 10 most enriched combinations of that particular spacing (**Fig. 5A-B**). Nucleotide composition of the spacer (as shown in **Fig. 5A-B**) was the mean nucleotide frequency across positions between both motif cores, relative to the corresponding nucleotide frequency between the same motif cores the input libraries. Preference for spacing (**Fig. S6B**) was computed as the change in the mean enrichment for the top 10 spaced combinations (i > 0) relative to the mean enrichment of the top 10 non-spaced combinations (i = 0, i.e. top 10 6mers): log2(enrichment_spaced_/enrichment_linear_). Significance was determined by setting a False Discovery Rate (FDR) using no-protein control libraries as follows: samples of 10 3mer cores were repeatedly drawn and the observed relative enrichments were used to set an FDR at each spacing *s*. Motif cores were sampled such that the relationships between sampled 3mers were the same as the relationship observed for that particular protein’s enriched cores.

### Assessment of flanking nucleotide context

For a given RBP, we only considered (protein-bound and input) reads that: a) contained one of the top 5 enriched 5mers; b) contained no additional secondary motifs, where secondary motifs were the top 50 enriched 5mers or all 5mers with an R-value >= 2, whichever set was larger. The remaining protein-bound and input reads were then subsampled to match the distribution of motifs and the positions of those motifs along a read. These reads were further subsampled to match the distribution of mean base pairing probabilities over the motif (bins used were [0-0.1),[0.1-0.2),…,[0.9,1.0]). For the analysis in **Fig. S6D**, protein-bound and input reads were instead subsetted only to reads where the motif was in a hairpin configuration (H5 in the MFE). The flanking nucleotide enrichment was then determined by centering these reads on the motif and computing the relative enrichment (log2(f_pulldown_NT / f_input_NT)) for each nucleotide at each position relative to the motif. We excluded the two nucleotides immediately adjacent to the motif on either side (to avoid capturing the extension of a core motif) as well as the first and last position of the random region in order to avoid certain nucleotide biases that can occur due to the presence of adaptor sequences. The overall enrichment (**Fig. 5G**) is the mean enrichment across all assessed positions, with significance assessed by a Wilcoxon rank-sum test.

Binding to mono-or dinucleotide rich sequence (**Fig. S6E**) in absence of a motif was done analogously, except only using reads that did not contain any of the top 50 5mers or 5mers with R >= 2. Enrichments for degenerate patterns were calculated as the mean of the 10 best degenerate *k*mers matching that pattern (e.g. mean of top 10/4096 12mers matching CCNNNCCNNNCC in the example in **Fig. 5H**, **S6H**). We first calculated enrichments for patterns where the fixed positions (e.g. CCCCCC in the previous example) contained only one or two nucleotides to assess which RBPs were biased towards binding to degenerate nucleotide-rich sequences, but later performed exhaustive searches where the fixed *k*mer was allowed to cover the entire sequence space (i.e. 4096 possible sequences in fixed positions × 210=(10 choose 4) patterns with 6 fixed positions and 6 internal Ns).

### Filter binding assay

Filter binding assay was performed with the oligo UUU(CCUCUCUUUUCC)UUU, i.e. the pattern CCNCNCNNNNCC flanked by Us and with Ns substituted with Us (in order to avoid creating high-affinity motifs and ensure the oligo was void of secondary structure). The negative control oligo used was U12. Custom RNA oligonucleotides were synthesized by IDT (Integrated DNA Technologies) and RBPs were purified as described earlier (see Cloning of RNA binding protein domains). RNA was end-labeled with ^32^P by incubating with Polynucleotide Kinase (NEB) according to manufacturer protocol. The assay was done following the protocol described in (Rio, 2012) for use with a 96-well dot-blot apparatus (Biorad). RBP and radio-labelled RNA were incubated in 50 uL binding buffer (500 uL 2M KCl, 10 uL 1M DTT, 400 uL 40% glycerol, 200 uL 1M Tris in 10 mL) for 1 hour at room temperature. Final concentration of RNA was 1nM and protein concentration ranged from 10nM-10uM (three-fold serial dilutions spanning this range).

### Calculation of feature-specific *R* values and relative entropy of context features

Feature-specific *R* values were calculated by assigning all 6mers into their respective bin for the feature under consideration for both the pulldown and input libraries, converting the counts into frequencies within each bin for both libraries, and computing the *R* value for the 6mer under consideration using the pulldown and input bin frequencies.

For **Fig. 6A-B**, bins used to compute feature-specific *R* values for each feature were the following: *P*_paired_: bin 1=0-0.2; bin 2=0.2-0.4; bin 3=0.4-0.6; bin 4=0.6-0.8; bin 5=0.8-1.0 *P*_flank_: bin 1=0-0.3; bin 2=0.3-0.45; bin 3=0.45-0.6; bin 4=0.6-0.75; bin 5=0.75-1.0 Core spacing: bin 1=0 nt spacing; bin 2 = 1 nt spacing; …; bin 11 = 10 nt spacing, where the spacing corresponds to the spacing between the two cores of a bipartite motif.

Nucleotide context: 16 bins, where the first four bins are quartiles of the percentage of A content flanking a 6mer based on the composition of input reads (bins 5-8, 9-12, and 13-16 are analogous for C, G, and U content, respectively). Each 6mer occurrence was therefore counted 4 times, into the corresponding bin for each of the four nucleotides.

Feature-specific *R* values within each bin were compared to the overall *R* value of the 6mer without binning (i.e. log2(*R*_bin_/*R*_original_)) to create the feature-specific enrichment profile for a particular context feature (example for *P*_paired_ for two RBPs in **Fig. S7A**).

For **Fig. 6C** and **Fig. S7C**, in order to compute the relative entropy of all context features on approximately the same scale, five bins as close to uniform in input as possible were created for each feature (if these five bins were exactly uniform, the maximum relative entropy would be log_2_(no. of bins) = log_2_(5) for each feature). The *P*_paired_ and *P*_flank_ bins were set individually for each RBP such that the five bins were equally populated for the 6mer under consideration for the given input library. Nucleotide context bins were created using the empirical distribution of nucleotide flanking contents for reads with the same 6mer in the input according to: bin 1) “high A” (flanking A content in the 75th percentile of input reads with the other three nucleotides each in their lower 50th percentile); bins 2), 3) and 4) “high C”, “high G”, and “high U” (analogous to high A for the respective nucleotides); bin 5) “other” (all nucleotides between the 30th and 50th percentile). Reads that did not fit into any of these categories were discarded. The five bins used for split motif spacing were: bin 1) spacing of 0 nt; bin 2) spacing of 1-2 nt; bin 3) 3-4 nt; bin 4) 5-7 nt; bin 5) 8-10 nt.

The relative entropy was then calculated for the probability distributions over the 5 bins for each of the four context features as:
DKL(f_6mer_PD ‖ f_6mer_IN), where D_KL_ is the Kullback-Leibler Divergence and f_6mer_PD and f_6mer_IN are length 5 vectors that sum to 1 as described by the bins above.

### Tissue specificity of RBP gene expression

Tissue specificity was measured as the information content deviation from a uniform distribution among all tissues as in (Gerstberger et al., 2014). For each RBP, the log_2_(TPM+1) was calculated for each of the 40 GTEx tissues (GTEx Consortium, 2013), and the tissue specificity was computed as the difference between the logarithm of the total number of samples (N=40) and the Shannon entropy of the expression values for an RBP:

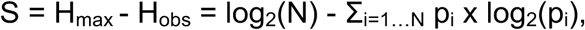

Where p_i_ = x_i_ / Σ_i=1…N_ X_i_

for x_i_ = log_2_(TPM_i_ + 1) in sample i.

The data used for the analyses were obtained from dbGaP accession number phs000424.v2.p1 in Jan. 2015. TPMs were measured using kallisto (Bray et al., 2016) on the following samples: Adipose-Subcutaneous: SRR1081567; AdrenalGland: SRR1120913; Artery-Tibial: SRR817094; Bladder: SRR1086236; Brain-Amygdala: SRR1085015; Brain-AnteriorCingulateCortex: SRR814989; Brain-CaudateBasalGanglia: SRR657731; Brain-CerebellarHemisphere: SRR1098519; Brain-Cerebellum: SRR627299; Brain-Cortex: SRR816770; Brain-FrontalCortex: SRR657777; Brain-Hippocampus: SRR614814; Brain-Hypothalamus: SRR661179; Brain-NucleusAccumben: SRR602808; Brain-SpinalCord: SRR613807; Brain-SubstantiaNigra: SRR662138; Breast-MammaryTissue: SRR1084674; Cervix: SRR1096057; Colon: SRR1091524; Esophagus: SRR1085211; FallopianTube: SRR1082520; Heart-LeftVentricle: SRR815517; Kidney-Cortex: SRR809943; Liver: SRR1090556; Lung: SRR1081283; MinorSalivaryGland: SRR1081589; Muscle-Skeletal: SRR820907; Nerve-Tibial: SRR612911; Ovary: SRR1102005; Pancreas: SRR1081259; Pituitary: SRR1077968; Prostate: SRR1099402; Skin: SRR807775; SmallIntestine: SRR1093314; Spleen: SRR1085087; Stomach: SRR814268; Testis: SRR1081449; Thyroid: SRR808886; Uterus: SRR820026; Vagina: SRR1095599.

